# Characterization of a mouse model of ICF syndrome reveals enhanced CD19 activation in inducing hypogammaglobulinemia

**DOI:** 10.1101/2023.03.09.531982

**Authors:** Zhengzhou Ying, Swanand Hardikar, Joshua B. Plummer, Tewfik Hamidi, Bin Liu, Yueping Chen, Jianjun Shen, Yunxiang Mu, Kevin M. McBride, Taiping Chen

## Abstract

Immunodeficiency, centromeric instability and facial anomalies (ICF) syndrome is a rare autosomal recessive disorder characterized by DNA hypomethylation and antibody deficiency. It is caused by mutations in *DNMT3B, ZBTB24, CDCA7* or *HELLS*. While progress has been made in elucidating the roles of these genes in regulating DNA methylation, little is known about the pathogenesis of the life-threatening hypogammaglobulinemia phenotype. Here we show that mice deficient for *Zbtb24* in the hematopoietic lineage recapitulate major clinical features of patients with ICF syndrome. Specifically, Vav-Cre-mediated ablation of *Zbtb24* does not affect lymphocyte development but results in reduced plasma cells and low levels of IgM, IgG1 and IgA. *Zbtb24*-deficient mice are hyper- and hypo-responsive to T-dependent and Tindependent type 2 antigens, respectively, and marginal zone B cell activation is impaired. B cells from *Zbtb24*-deficient mice display elevated CD19 phosphorylation. Heterozygous disruption of *Cd19* can revert the hypogammaglobulinemia phenotype in these mice. Mechanistically, *Il5ra* (interleukin-5 receptor subunit alpha) is derepressed in *Zbtb24*-deficient B cells, and elevated IL-5 signaling enhances CD19 phosphorylation. Our results reveal a novel link between IL-5 signaling and CD19 activation and suggest that abnormal CD19 activity contributes to immunodeficiency in ICF syndrome.

**SIGNIFICANCE STATEMENT:** ICF syndrome is a rare immunodeficiency disorder first reported in the 1970s. The lack of appropriate animal models has hindered the investigation of the pathogenesis of antibody deficiency, the major cause of death in ICF syndrome. Here we show that, in mice, disruption of *Zbtb24*, one of the ICF-related genes, in the hematopoietic lineage results in low levels of immunoglobulins. Characterization of these mice reveals abnormal B cell activation due to elevated CD19 phosphorylation. Mechanistically, *Il5ra* (interleukin-5 receptor subunit alpha) is derepressed in *Zbtb24*-deficient B cells, and increased IL-5 signaling enhances CD19 phosphorylation.

## INTRODUCTION

Immunodeficiency, centromeric instability and facial anomalies (ICF) syndrome is a rare autosomal recessive disorder characterized by antibody deficiency, facial dysmorphism, failure to thrive, and mental retardation (1, 2). A cytogenetic feature of ICF syndrome is chromosomal rearrangements due to DNA hypomethylation of satellite repeats at (peri)centromeric heterochromatin (1, 3–6). Four genes have been identified to be mutated in ICF syndrome – *DNMT3B* (ICF1), *ZBTB24* (ICF2), *CDCA7* (ICF3) and *HELLS* (ICF4) – with ICF1 (~50%) and ICF2 (~30%) accounting for the vast majority of cases (7–13). Recent evidence suggests the four proteins to be components of a pathway that regulates the specificity of DNA methylation. Specifically, ZBTB24, a member of the zinc-finger (ZF)- and BTB domain-containing (ZBTB) family of transcriptional regulators, induces *CDCA7* expression, and CDCA7 recruits HELLS, a DNA helicase involved in chromatin remodeling, to (peri)centromeric heterochromatin to facilitate DNA methylation by the DNA methylation machinery, including DNMT3B (14–19).

Patients with ICF syndrome suffer from recurrent and often fatal infections in early childhood due to hypogammaglobulinemia or agammaglobulinemia. While they usually have normal counts of T and B lymphocytes, phenotypic analysis revealed the presence of only naïve B cells and the absence of memory B cells and plasma cells, suggesting impaired B cell activation and/or terminal differentiation (20). However, the cellular and molecular defects underlying antibody deficiency in ICF syndrome remain poorly understood.

B cell survival and differentiation are tightly controlled by signaling from the B cell receptor (BCR), a transmembrane complex composed of a membrane-bound immunoglobulin (Ig) and the signal transduction components Igα/Igβ (CD79a/CD79b). The BCR transmits constitutive, tonic signals necessary for survival in the absence of antigen binding and activation signals following antigen engagement (21). For mature B cells in the periphery, BCR engagement activates signaling pathways that induce proliferation and transcriptional programs for differentiation into germinal centers (GCs), memory and plasma cells (22). However, during early B cell development, BCR activation plays a key role in central tolerance. BCR engagement in bone marrow (BM) immature cells facilitates elimination of self-reactive BCRs (23). Thus, BCR activation controls B cell fate in a context- and developmental stage-dependent manner.

CD19, a transmembrane protein expressed throughout B cell development, modulates BCR signaling. CD19-deficient (*Cd19^-/-^*) mice develop B cells but have reduced peripheral populations and deficient GC, B1 and marginal zone (MZ) compartments (24). B cells without CD19 are hyporesponsive to BCR engagement with reduced phosphorylation of signaling factors and reduced proliferation, and loss of terminal differentiation (25). CD19 overexpression results in the opposite phenotype, as B cells from human CD19-transgenic (hCD19TG) mice display enhanced BCR signaling and elevated antibody response. hCD19TG mice also have reduced levels of naïve cells leaving the BM, presumably due to overactive negative selection (26). Further characterization of the *Cd19^-/-^* and hCD19TG mice revealed that CD19 enhances humoral immune responses to T-dependent (TD) antigens but inhibits humoral immune response to T-independent type 2 (TI-2) antigens (27). Thus, CD19 has been proposed to function as a critical rheostat that amplifies BCR signaling and lowers thresholds for activation (28). While the relationship between BCR and CD19 signaling is well established, the full catalog of factors that influence this signaling axis remain to be defined.

In this study, we show that conditional ablation of *Zbtb24* in the hematopoietic lineage in mice results in a phenotype that mimics major clinical manifestations of ICF syndrome, including normal T and B cell development, reduced plasma cells, and hypogammaglobulinemia. We provide evidence that Zbtb24 is a negative regulator of CD19 activity, as *Zbtb24*-deficient B cells display elevated CD19 phosphorylation, which contributes to hypogammaglobulinemia. Mechanistically, *Il5ra*, encoding the IL-5-binding subunit (α chain) of the IL-5 receptor (IL-5Rα), is derepressed in *Zbtb24*-deficient B cells, and the resulting increase in IL-5 signaling enhances CD19 phosphorylation.

## RESULTS

### Mice deficient for *Zbtb24* in hematopoietic cells recapitulate the antibody deficiency phenotype of ICF syndrome

To investigate the biological functions of Zbtb24, we generated a *Zbtb24* conditional allele (*Zbtb24^2lox^*) in mice, which contains two *loxP* sites flanking a 1.3-kb region including exon 2. Cre-mediated deletion of the floxed region removes the N-terminal 316 amino acids that include the BTB domain, AT-hook and part of the ZF domain, resulting in a null allele (*Zbtb24^-^*) (Figure S1A). Genotypes were determined by Southern blotting and PCR (Figure S1B-D). *Zbtb24^2lox/2lox^*, *Zbtb24^2lox/-^* and *Zbtb24^+/-^* mice were all grossly normal and fertile. However, intercrosses between *Zbtb24^+/-^* or *Zbtb24^2lox/-^* mice produced no viable pups with homozygous deletion (*Zbtb24^-/-^*). Timed mating revealed that *Zbtb24^-/-^* embryos were arrested before 8.5 days post coitum (Figure S1E), in agreement with a previous report (14). Thus, *Zbtb24* is essential for embryonic development in mice, although ICF2 patients – many carrying homozygous or compound heterozygous null mutations in *ZBTB24* (11, 12, 29, 30) – are viable.

To assess the role of Zbtb24 in immune cells, we conditionally deleted *Zbtb24* in the hematopoietic lineage using Vav-Cre (31). PCR genotyping and Western blotting confirmed efficient deletion of *Zbtb24* in B220+ B cells and CD3+ T cells, but not in tail samples (Figure 1A, B). *VavCre:Zbtb24^2lox/2lox^* (VAV-CKO) mice grew normally, were fertile, and appeared indistinguishable from *VavCre:Zbtb24^2lox/+^* (control) and wild-type (WT) mice.

**Figure 1.**
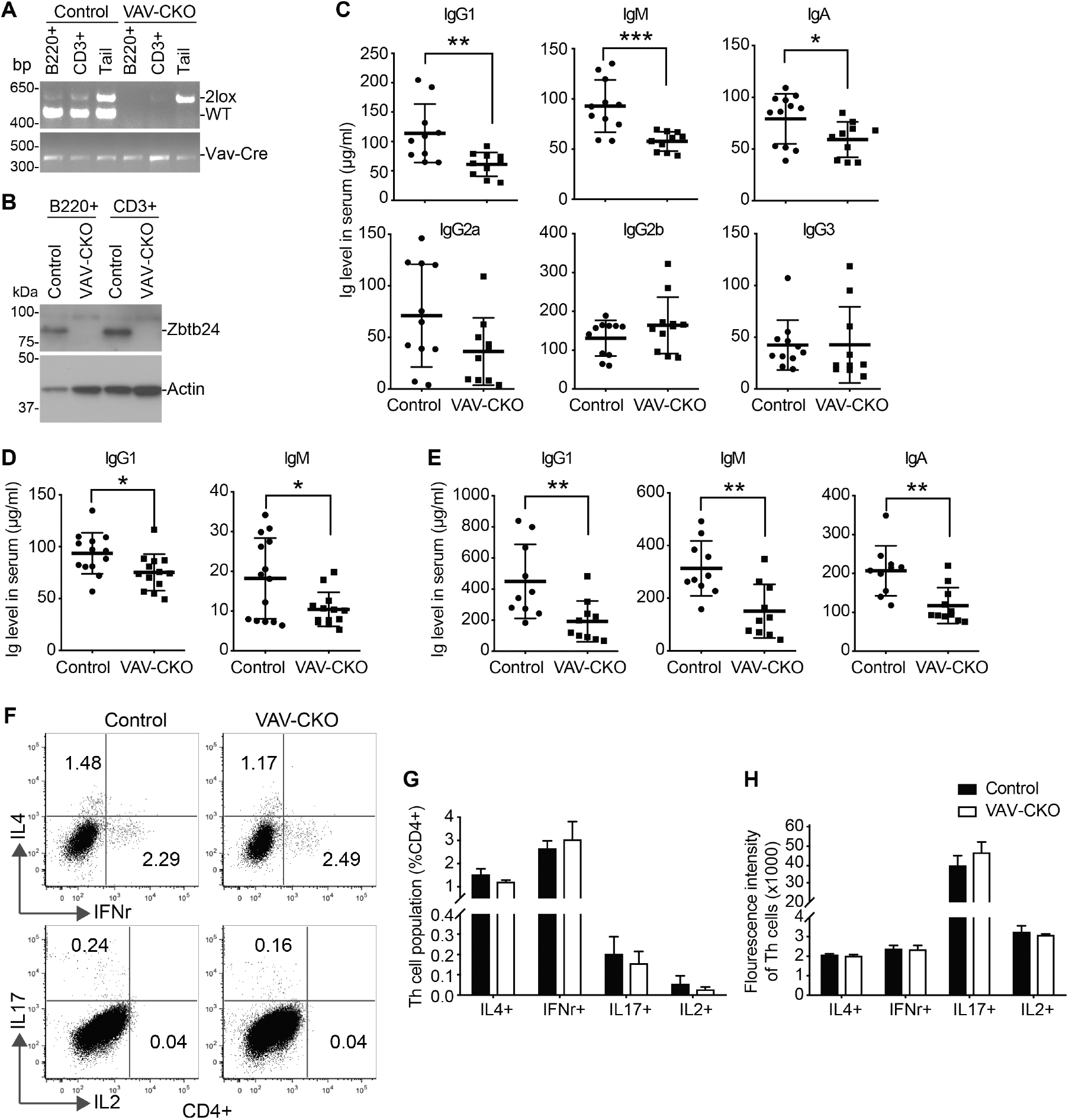
VAV-Cre-mediated *Zbtb24* ablation results in hypogammaglobulinemia in mice. (A) PCR genotyping of sorted B220+ B cells and CD3+ T cells and tail DNA from Control (*VavCre:Zbtb24^2lox/+^*) and VAV-CKO (*VavCre:Zbtb24^2lox/2lox^*) mice. (B) Western blot showing *Zbtb24* depletion in sorted B220+ B cells and CD3+ T cells from VAV-CKO mice. (C-E) ELISA analysis of sera from Control and VAV-CKO mice showing basal levels of IgG1, IgM, IgA, IgG2a, IgG2b and IgG3 at 8-10 weeks of age (C), IgG1 and IgM at 3 weeks of age (D) and IgG1, IgM and IgA at 6 months of age (E). (F-H) Analysis of IL4-, IFNγ-, IL17-, and IL2-producing CD4+ cells in spleens from Control and VAV-CKO mice. Shown are representative flow cytometry data (F), percentages of Th subpopulation in CD4+ cells (G) and fluorescence intensity of the Th cells (H). Presented are data (mean±SD) from 10-14 (C-E) and 8 (G, H) mice, respectively. Statistical analysis was done using Student’s t-test. *P<0.05; **P<0.01; ***P<0.001.

A major clinical feature of ICF syndrome is hypogammaglobulinemia (32). We measured the basal serum Ig levels in VAV-CKO mice. Compared to control littermates, adult (8-10-week-old) VAV-CKO mice had significantly lower levels of IgG1, IgM and IgA, slightly lower levels of IgG2a, and comparable levels of IgG2b and IgG3 (Figure 1C). Hypogammaglobulinemia was also detected in younger (3-week-old) and older (6-month-old) VAV-CKO mice (Figure 1D, E). With the increasing Ig levels over time, the differences between control and *Zbtb24*-deficient mice seemed to become more evident (Figure 1C-E). Control and VAV-CKO mice showed no differences in cytokine secretion by helper T cells, including IFNγ, IL-4, IL-2, and IL-17 (Figure 1F-H), suggesting that hypogammaglobulinemia induced by *Zbtb24* deficiency was caused by B cell-intrinsic defects. Consistent with the notion, Lck-Cre-mediated *Zbtb24* deletion in the T cell lineage had no effects on basal serum Ig levels, T cell subtypes in spleen, and humoral immune response to the TD antigen nitrophenyl-conjugated Keyhole Limpet Hemocyanin (NP-KLH) (Figure S2A-C).

### VAV-CKO mice show no defects in lymphocyte development but have decreased plasma cells

To explore the cellular defects underlying hypogammaglobulinemia in VAV-CKO mice, we examined lymphocyte subtypes. Flow cytometry showed that *Zbtb24*-deficient mice had normal percentages of B lineage (B220+), T lineage (CD3+), and myeloid lineage (CD11b+) cells in BM (Figure S3A, B) and normal total numbers of cells in BM and spleen (Figure S3C, D). Analysis of pro-B, pre-B, immature and mature B cells in BM indicated that B cell development progressed appropriately in the absence of Zbtb24 (Figure 2A, Figure S3E). Early subtypes of T cell development in thymus were also normal in VAV-CKO mice (Figure S3F, G). In addition, *Zbtb24* deficiency had no effect on the subpopulations of naïve B cells in spleen, including follicular (FO), MZ, and transitional B cells (Figure 2B, Figure S3H). Similarly, the frequencies of naïve, effector, and memory T cell subtypes in spleen showed no differences between VAV-CKO and control littermates (Figure 2C, Figure S3I). However, there were significant decreases in the frequencies and numbers of plasma cells in the BM and spleen of *Zbtb24*-deficient mice (Figure 2D-G). These results, consistent with the observation that ICF patients have normal B and T cell counts (33, 34), suggest that Zbtb24 is not required for lymphocyte development but plays a role in B cell activation and/or terminal differentiation.

**Figure 2.**
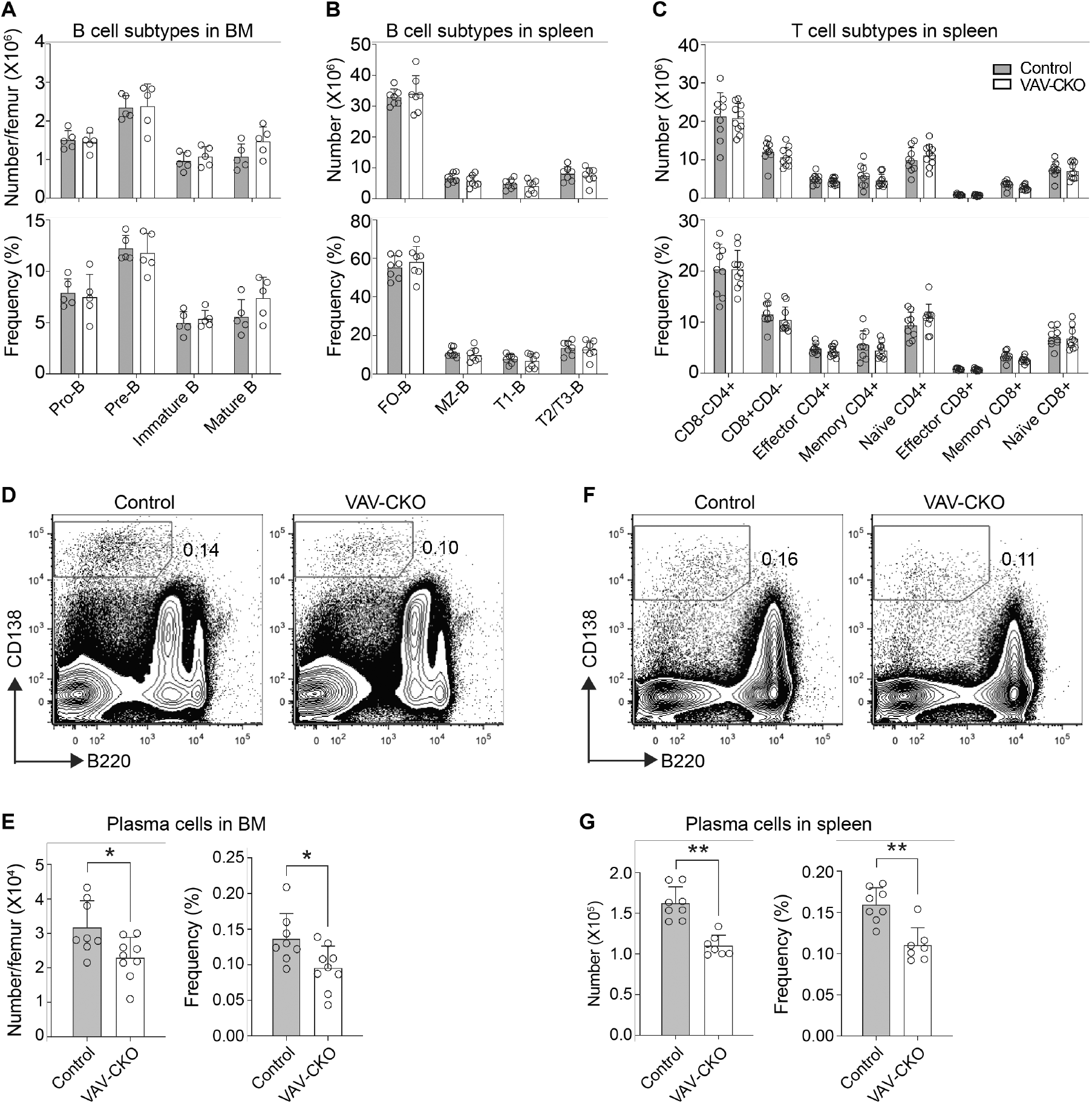
*Zbtb24* deficiency has no effect on B and T cell development but leads to decreased plasma cells. (A-C) Numbers (top) and frequencies (bottom) of B cells at various maturation stages [pro-B (B220+CD43+), pre-B (B220+CD43-IgM-IgD-), immature (B220+CD43-IgM+IgD-), and mature (B220+CD43-IgM+IgD+) B cells] in BM (A), various B cell subtypes [FO (B220+CD93-CD23+CD21/35lo), MZ (B220+CD93-CD23-CD21/35hi), T1 (B220+CD93+CD23-), and T2/T3 (B220+CD93+CD23+) B cells] in spleen (B) and various T cell subtypes [helper (CD8-CD4+), cytotoxic (CD8+CD4-), effector helper (CD4+CD8-CD44+CD62L-), memory helper (CD4+CD8-CD44+CD62L+), naïve helper (CD4+CD8-CD44-CD62L+), effector cytotoxic (CD4-CD8+CD44+CD62L-), memory cytotoxic (CD4-CD8+CD44+CD62L+), and naïve cytotoxic (CD4-CD8+CD44-CD62L+) T cells] in spleen (C) from Control and VAV-CKO mice. (D-G) Flow cytometry analysis of plasma cells (CD11b-CD3-B220lo/CD138hi) in BM (D, E) and spleen (F, G) from Control and VAV-CKO mice. Shown are representative flow cytometry data (D, F) and absolute numbers (left) and frequencies (right) of plasma cells (E, G). Presented in (A, B, C, E, G) are data (mean±SD) from 5-9 mice. Statistical analysis was done using Student’s t-test. *P<0.05; **P<0.01.

### VAV-CKO mice show divergent humoral immune responses to T-dependent and -independent antigens

To assess the impact of *Zbtb24* deficiency on lymphocyte functions, we immunized mice with TD and TI antigens. Flow cytometry revealed that unimmunized (day 0) control and VAV-CKO mice had similarly low percentages and numbers of GC (GL7+Fas+) B cells in their spleens. Fourteen days after immunization with NP-KLH, a TD antigen, all mice had significantly higher numbers of GC B cells in their spleens, but the increases were consistently greater in VAV-CKO mice (>10 fold) than in control mice (~5 fold) (Figure 3A, B, Figure S4A). Immunofluorescence staining of spleen sections with peanut agglutinin (PNA) also showed increases in the numbers and sizes of GCs in *Zbtb24*-deficient mice (Figure S4B, C). As measured by ELISA in sera collected 14, 28 and 56 days after NP-KLH injection, steady increases of low-affinity (anti-NP20) and high-affinity (anti-NP7) IgG1 against NP and transient increases of low- and high-affinity IgM against NP were observed in both control and VAV-CKO mice, with significantly higher titers in *Zbtb24*-deficient mice (Figure 3C, D). IgA and IgG2a were undetectable at day 14 after immunization, and IgG2b and IgG3 responses were comparable in control and VAV-CKO mice at all time points examined (Figure S4D). In sera collected 14 days after the secondary immunization (administered 60 days after the primary immunization), significantly higher titers of low-affinity and comparable levels of high-affinity anti-NP specific IgG1 and IgM were detected in VAV-CKO mice, as compared to control littermates (Figure 3E, F). Based on the ratios of high-affinity to low-affinity antibodies (Figure 3G, H), VAV-CKO mice were apt to produce low-affinity IgG1 and IgM. Of note, previous work reported that mice deficient for *Dnmt3b* (another ICF-related gene) and its paralog *Dnmt3a* in B cells also show normal B cell development but exhibit enhanced TD responses upon immunization (35).

**Figure 3.**
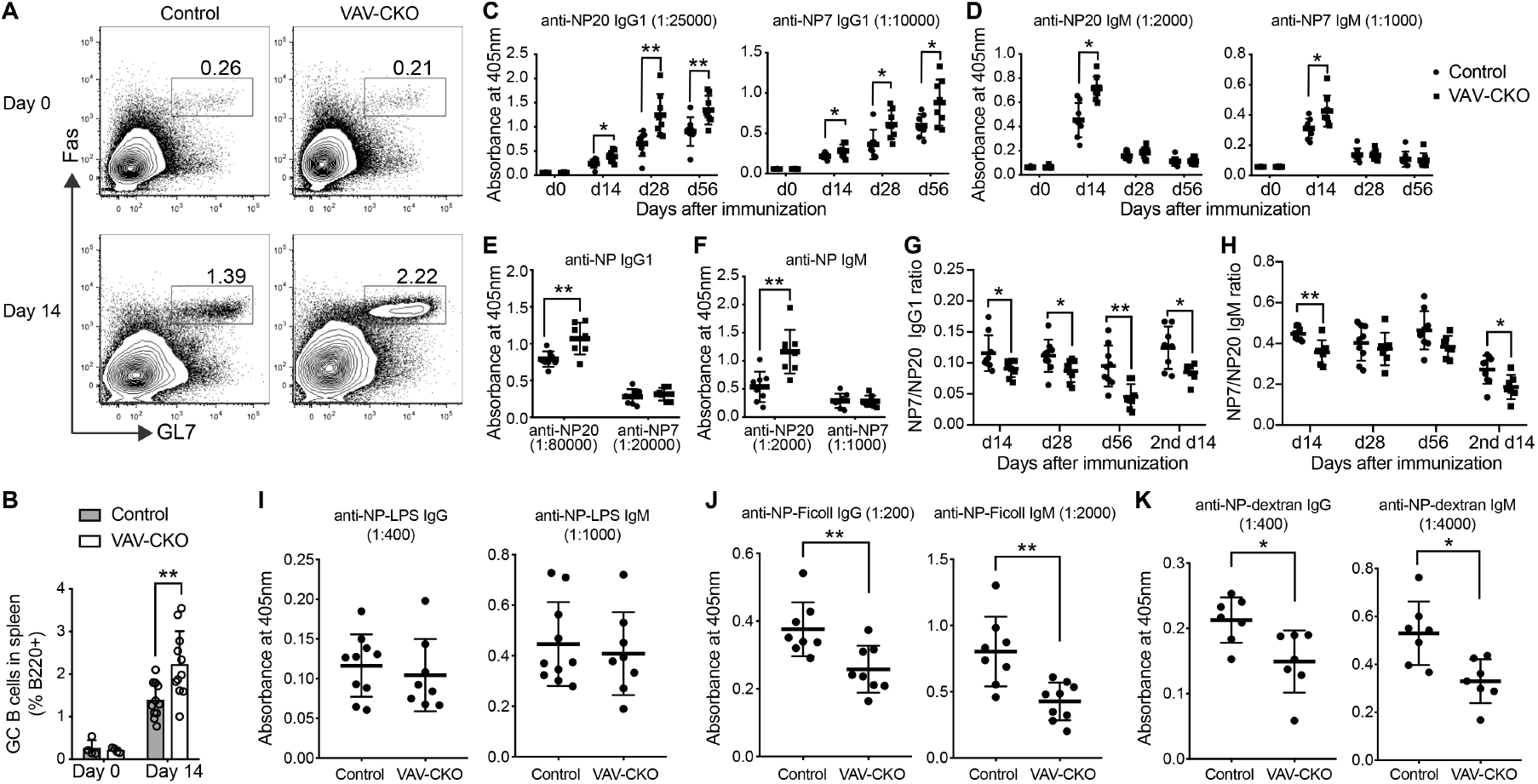
*Zbtb24*-deficient mice show different humoral immune responses to T-dependent and T-independent antigens. (A, B) Flow cytometry analysis of germinal center (GC) B cells (B220+GL7+Fas+) before (day 0) and 14 days after immunization with NP-KLH in spleens from Control and VAV-CKO mice. Shown are representative flow cytometry data (A) and frequencies of GC B cells (percentages in B220+ cells) in spleen (B). (C-F) Relatively low-affinity (NP20) and high-affinity (NP7) anti-NP specific IgG1 (C, E) and IgM (D, F) antibody absorbance by ELISA in sera from Control and VAV-CKO mice at day 0, day 14, day 28, and day 56 after first injection with NP-KLH (C, D) and at day 14 after second injection with NP-KLH (E, F). (G, H) Ratio of high-affinity to low-affinity IgG1 (G) and IgM (H) in Control and VAV-CKO mice at day 0, day 14, day 28, and day 56 after first injection with NP-KLH and at day 14 after second injection with NP-KLH. (IK) Anti-NP specific IgG and IgM antibody absorbance by ELISA in sera from Control and VAV-CKO mice at day 6 after immunization with NP-LPS (I), NP-Ficoll (J), or NP-dextran (K). Presented in (B-K) are data (mean±SD) from 8-10 mice. Statistical analysis was done using Student’s t-test. *P<0.05; **P<0.01.

While hyperresponsive to TD antigens, *Zbtb24*-deficient mice had lower natural antibodies (Figure 1C-E), suggesting that immune responses to T-independent antigens may also be perturbed by loss of Zbtb24. There are two categories of TI antigens: TI-1 antigens have the ability to directly activate B cells, and TI-2 antigens are typically repeating polymers and cause simultaneous crosslinking of BCRs on B lymphocytes. After immunization with NP-LPS, a TI-1 antigen, control and VAV-CKO mice had similar titers of IgG and IgM against NP in their sera (Figure 3I). However, after immunization with NP-Ficoll or NP-Dextran, both being TI-2 antigens, VAV-CKO mice mounted significantly lower anti-NP specific IgG and IgM responses than control littermates (Figure 3J, K).

In summary, our results show that *Zbtb24* deficiency results in enhanced and diminished humoral immune responses to TD and TI-2 antigens, respectively.

### *Zbtb24*-deficient marginal zone B cells are defective in proliferation in response to stimulation with anti-IgM

To gain insights into the intrinsic function of Zbtb24 in B cells, we tested the ability of B cells to be activated in the absence of T cell help. FO and MZ B cells sorted from control and VAV-CKO mice were stained with CellTrace Violet (CTV) reagent, then stimulated with LPS or soluble F(ab’)2 fragment of anti-IgM (referred to as anti-IgM hereafter), an analogue of TI-2 antigen, for 72 hours and analyzed with flow cytometry to monitor cell proliferation by dye dilution. After stimulation with LPS, both FO and MZ B cells from VAV-CKO mice were activated and proliferated normally (Figure 4A, B). After stimulation with anti-IgM, whereas the vast majority of FO B cells from VAV-CKO mice underwent proliferation, similar to those from control mice, significantly smaller fractions of *Zbtb24*-deficient MZ B cells underwent proliferation when compared to their control counterparts (Figure 4C, D).

**Figure 4.**
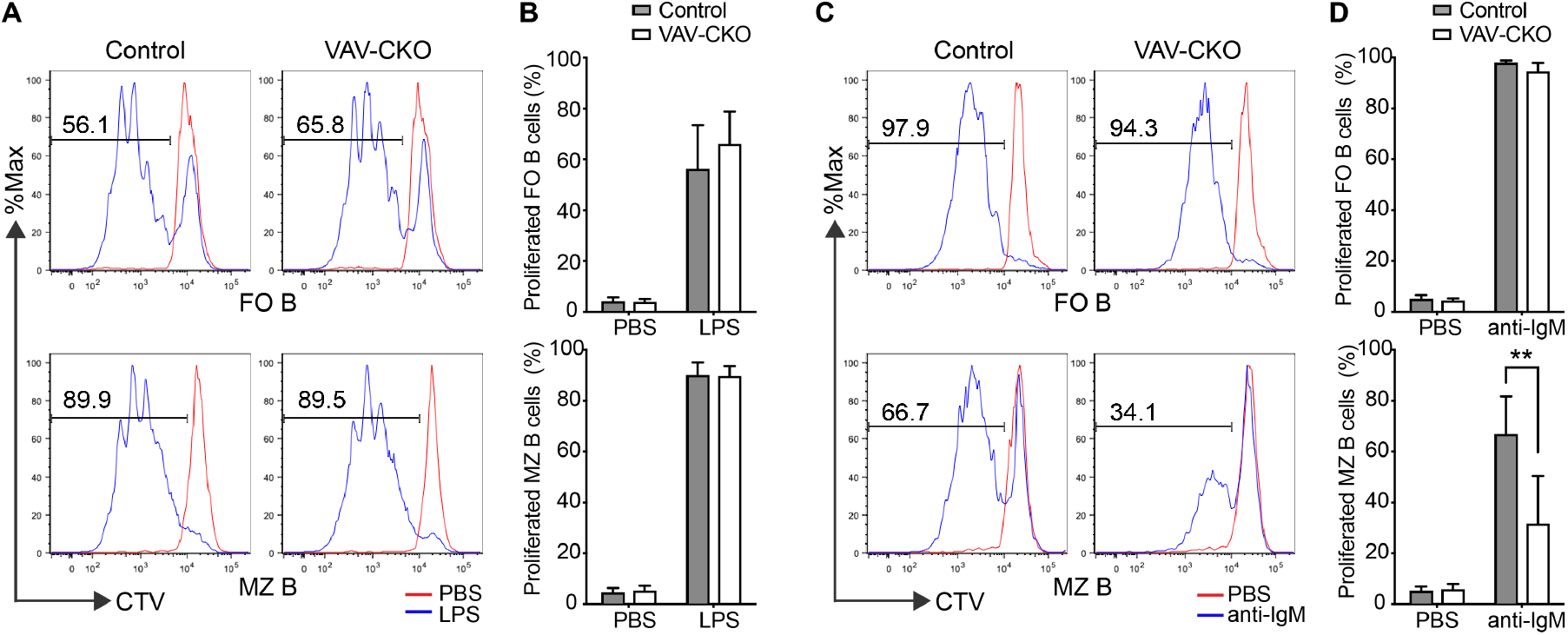
*Zbtb24*-deficient MZ B cell activation is impaired after stimulation with anti-IgM. CellTrace Violet (CTV) assay showing proliferation of FO and MZ B cells from Control and VAV-CKO mice after 72 hours of stimulation with LPS (A, B) or anti-IgM (C, D), with cells treated with PBS serving as unstimulated control. Shown are representative flow cytometry data (A, C) and percentages (mean±SD from 8 mice per genotype) of proliferated FO and MZ B cells (B, D). Statistical analysis was done using Student’s t-test. **P<0.01.

### Enhanced CD19 activity contributes to hypogammaglobulinemia in VAV-CKO mice

Our results suggested that antibody deficiency in VAV-CKO mice was a B cell-intrinsic phenotype (Figures 1 and 4, Figure S2). To further confirm that, we deleted *Zbtb24* in the B cell lineage using CD19-Cre (36). To our surprise, *Cd19Cre:Zbtb24^2lox/2lox^* (CD19-CKO) mice had normal levels of serum IgG1, IgM and IgA (Figure 5A), although Zbtb24 depletion was robust in B cells (Figure S5A). These mice also responded normally to immunization with NP-KLH, as judged by the frequencies of total GC B cells and anti-NP specific IgG1 and IgM GC and memory B cells in spleen (Figure S5B-D). Given that one copy of *Cd19* is disrupted in the CD19-Cre knock-in (KI) mouse (36), it is possible that different levels of CD19 expression contributed to the phenotypic differences between CD19-CKO and VAV-CKO mice. Indeed, the divergent humoral immune responses of VAV-CKO mice to different antigens – i.e. hyperresponsive to TD and hyporesponsive to TI-2 antigens (Figure 3) – were similar to the phenotype of hCD19TG mice and opposite to that of *Cd19^-/-^* mice (27). We therefore asked if *Cd19* expression and/or function were altered in B cells from VAV-CKO mice. While the *Cd19* transcript and protein levels were normal in *Zbtb24*-deficient FO and MZ B cells, the levels of CD19 phosphorylation in these cells were substantially higher than those in control cells, both before and after stimulation with anti-IgM (Figure 5B, C). By contrast, B cells from CD19-CKO showed a ~50% reduction in CD19 expression (as expected) and similar levels of CD19 phosphorylation after stimulation with anti-IgM, as compared to B cells from WT mice (Figure 5D, Figure S5E). Based on these results, we hypothesized that enhanced CD19 activity contributed to the phenotype of VAV-CKO mice.

**Figure 5.**
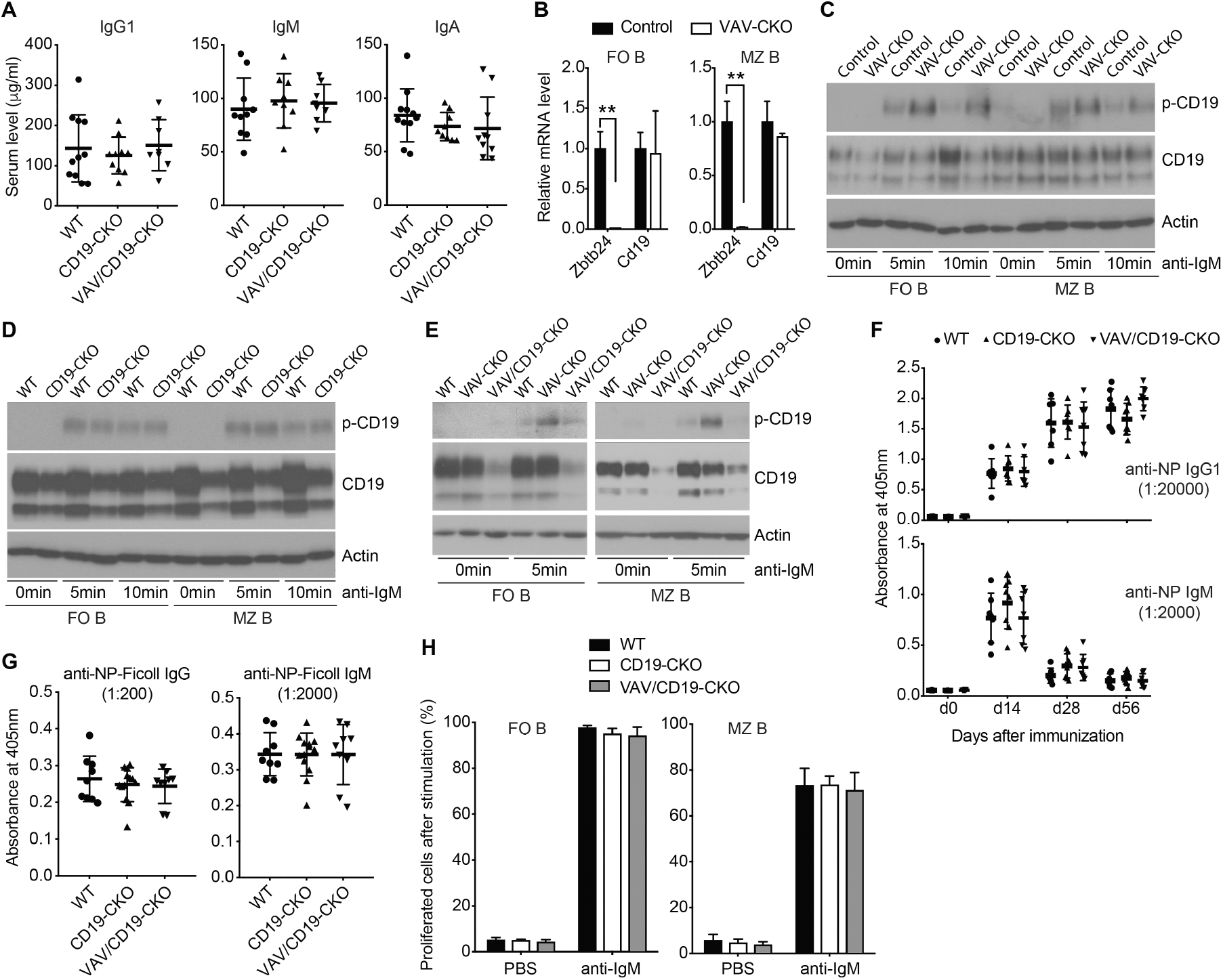
The immune phenotype of VAV-CKO mice is largely due to enhanced CD19 phosphorylation. (A) Basal IgG1, IgM and IgA levels in sera from 8-10-week-old WT, CD19-CKO (*Cd19Cre:Zbtb24^2lox/2lox^*) and VAV/CD19-CKO (*VavCre:Cd19Cre:Zbtb24^2lox/2lox^*) mice. (B) RT-qPCR data (mean±SD from 3 mice per genotype) showing relative mRNA levels of *Zbtb24* and *Cd19* in sorted FO and MZ B cells from Control and VAV-CKO mice. Statistical analysis was done using Student’s t-test. **P<0.01. (C-E) Western blots showing the levels of phosphorylated CD19 (p-CD19) and total CD19 in sorted FO and MZ B cells from Control and VAV-CKO mice (C), WT and CD19-CKO mice (D) or WT, VAV-CKO and VAV/CD19-CKO mice (E) that were unstimulated (0 min) or stimulated with anti-IgM for 5 or 10 min. (F) Anti-NP specific IgG1 and IgM antibody absorbance by ELISA in sera from WT, CD19-CKO and VAV/CD19-CKO mice at day 0, day 14, day 28, and day 56 after immunization with NP-KLH. (G) Anti-NP specific IgG and IgM antibody absorbance by ELISA in sera from WT, CD19-CKO and VAV/CD19-CKO mice at day 6 after immunization with NP-Ficoll. (H) Percentages of proliferated FO and MZ B cells from WT, CD19-CKO and VAV/CD19-CKO mice after 72 hours of stimulation with PBS or anti-IgM by CTV assay.

To test the hypothesis, we generated VAV-CKO mice that were heterozygous for *Cd19* by introducing the CD19-Cre KI allele through breeding. In contrast to hypogammaglobulinemia in VAV-CKO mice (Figure 1), *VavCre:Cd19Cre:Zbtb24^2lox/2lox^* (VAV/CD19-CKO) mice had normal levels of serum IgG1, IgM and IgA (Figure 5A). Similarly, B cells from VAV/CD19-CKO mice no longer exhibited hyperphosphorylation of CD19 in response to stimulation with anti-IgM (Figure 5E). VAV/CD19-CKO mice, as well as CD19-CKO mice, also showed normal humoral immune responses to immunization with NP-KLH (Figure 5F) and NP-Ficoll (Figure 5G). Consistent with the in vivo data, sorted FO and MZ B cells from VAV/CD19-CKO and CD19-CKO mice proliferated normally after stimulation with anti-IgM (Figure 5H), unlike MZ B cells from VAV-CKO mice that were defective in proliferation (Figure 4C, D). Taken together, these results demonstrate that the immune phenotype of VAV-CKO mice can be rescued by reducing *Cd19* expression, supporting our hypothesis.

### Increased IL-5 signaling contributes to enhanced CD19 phosphorylation in *Zbtb24*-deficient B cells

To explore the molecular mechanism by which *Zbtb24* deficiency leads to enhanced CD19 phosphorylation, we performed RNA sequencing (RNA-Seq) using sorted naïve FO and MZ B cells from control and VAV-CKO mice. Bioinformatics analysis, using adjusted pvalue < 0.05 as the significance cutoff, identified only small numbers of differentially expressed genes in *Zbtb24*-deficient samples – 43 genes (21 downregulated and 22 upregulated) in FO B cells and 35 genes (11 downregulated and 24 upregulated) in MZ B cells – with 24 common genes (8 downregulated and 16 upregulated) in both FO and MZ B cells (Figure 6A). Some of the common downregulated genes, including *Cdca7, Taf6, Cdc40* and *Ostc*, have been identified as direct target genes of Zbtb24 in other cell types (16, 17). Interestingly, among the common upregulated genes were *Crisp3* (cysteine-rich secretory protein 3) and *Il5ra* (interleukin 5 receptor subunit alpha), which have been implicated in B cell functions (37–40). RT-qPCR analysis verified the downregulation of *Cdca7* and upregulation of *Crisp3* and *Il5ra* in *Zbtb24*-deficient FO and MZ B cells (Figure 6B).

**Figure 6.**
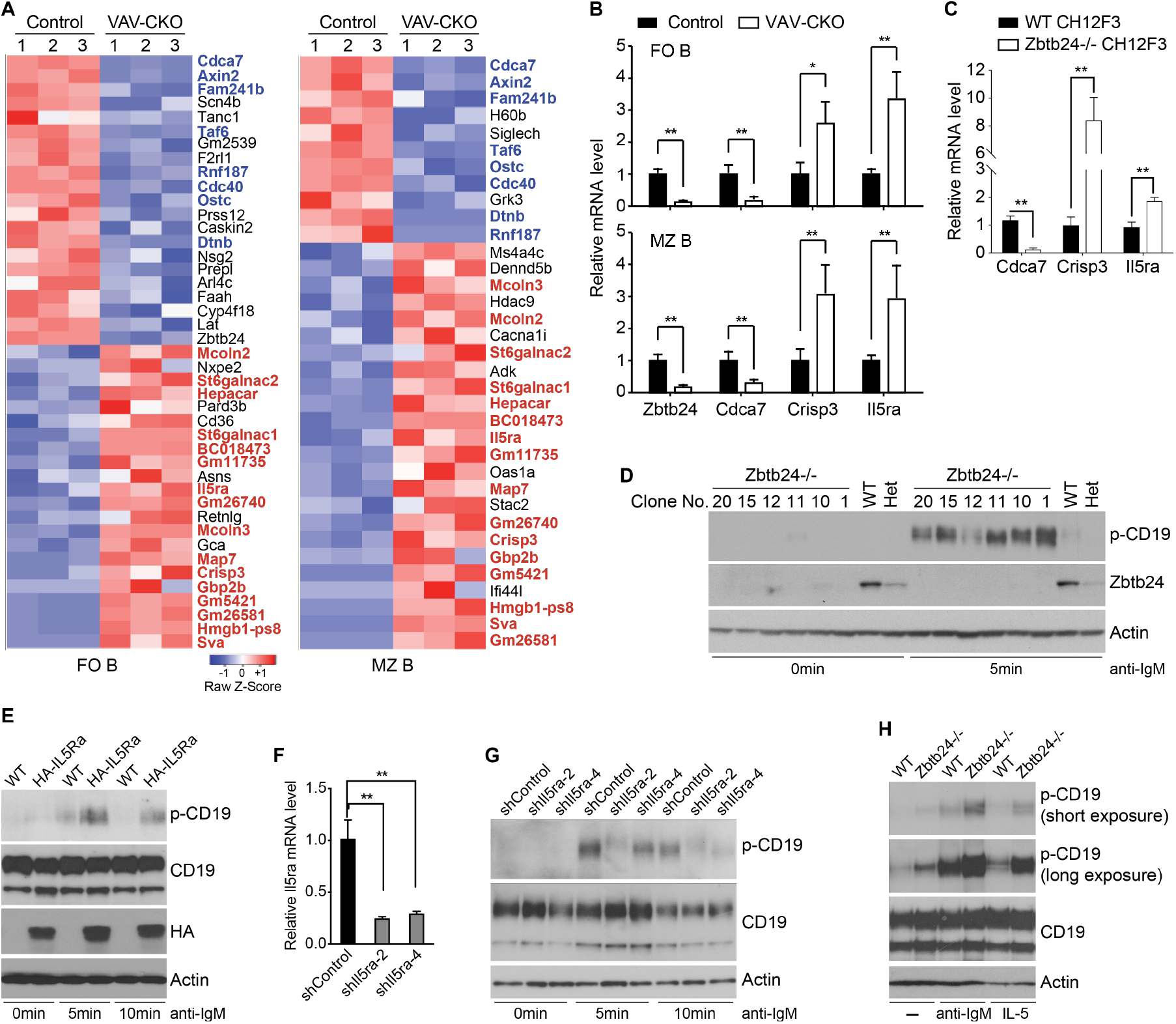
Increased IL-5 signaling contributes to enhanced CD19 phosphorylation in *Zbtb24-* deficient B cells. (A) Heatmap of differentially expressed genes in naïve FO and MZ B cells from Control and VAV-CKO mice identified by RNA-Seq. Genes that are downregulated and upregulated in both FO and MZ B cells are highlighted in blue and red, respectively. (B, C) RT-qPCR analysis verifying downregulation of *Cdca7* and upregulation of *Crisp3* and *Il5ra* in *Zbtb24-deficient* FO and MZ B cells (B) and *Zbtb24^-/-^* CH12F3 cells (C). (D) Western blots showing that, after stimulation with IgM, CD19 phosphorylation is dramatically enhanced in *Zbtb24^-/-^* CH12F3 cells, compared to WT and *Zbtb24*^+/-^ (Het) CH12F3 cells. (E) Overexpression of HA-tagged IL-5Rα in CH12F3 cells recapitulates the effect of *Zbtb24* deficiency on CD19 phosphorylation. (F, G) shRNA-mediated *Il5ra* knockdown in *Zbtb24^-/-^* CH12F3 cells reduces CD19 phosphorylation in response to anti-IgM stimulation. Shown are RT-qPCR data demonstrating the knockdown effects of the two most efficient *Il5ra* shRNAs, i.e. *shIl5ra-2* and *shIl5ra-4* (F) and Western blots with the indicated antibodies (G). (H) Western blots showing that IL-5 treatment (10 ng/ml for 5 min) induces enhanced CD19 phosphorylation in *Zbtb24^-/-^* CH12F3 cells, similar to the effect of anti-IgM. Statistical analysis was done using Student’s t-test. *P<0.05; **P<0.01.

In an attempt to find an appropriate cell line for validation experiments, we used CRISPR/Cas9 gene editing to disrupt *Zbtb24* in CH12F3 cells, a murine B cell lymphoma cell line that shows abundant expression of Zbtb24 and can be activated by anti-IgM (19, 41). Multiple clones with frameshift indels on both copies of *Zbtb24* were obtained (Figure S6A, B). *Zbtb24* ablation in CH12F3 cells resulted in gene expression changes that were similar to those in *Zbtb24*-deficient naïve B cells, including downregulation of *Cdca7* and upregulation of *Crisp3* and *Il5ra* (Figure 6C). *Zbtb24^-/-^* CH12F3 cells also showed dramatic increases in CD19 phosphorylation after stimulation with anti-IgM, as compared to WT and *Zbtb24^+/-^* CH12F3 cells (Figure 6D).

We then investigated the relevance of *Crisp3* and *Il5ra* upregulation in enhancing CD19 phosphorylation using CH12F3 cells. While overexpression of HA-tagged Crisp3 showed no obvious effect on CD19 phosphorylation (Figure S6C), overexpression of HA-IL-5Rα recapitulated the effect of *Zbtb24* deficiency – i.e. enhanced CD19 phosphorylation after stimulation with anti-IgM (Figure 6E, F). Conversely, shRNA-mediated *Il5ra* knockdown (KD) in *Zbtb24^-/-^* CH12F3 cells substantially prevented hyperphosphorylation of CD19 in response to anti-IgM stimulation (Figure 6G, H). *Il5ra* encodes the IL-5-binding subunit of IL-5 receptor. We therefore tested the possibility that enhanced CD19 phosphorylation in *Zbtb24*-deficient B cells was due to increased IL-5 signaling. Indeed, IL-5 treatment (10 ng/ml for 5 min) induced markedly higher CD19 phosphorylation in *Zbtb24^-/-^* CH12F3 cells than in WT CH12F3 cells (Figure 6I). Collectively, these results indicate that Zbtb24 is a negative regulator of *Il5ra* expression in B cells and that, in the absence of Zbtb24, elevated IL-5 signaling, as a result of derepression of *Il5a*, contributes to enhanced CD19 activity.

## DISCUSSION

ICF syndrome was first reported in the 1970s (42–44). Yet, little progress has been made in understanding the pathogenesis of the life-threatening hypogammaglobulinemia phenotype, partly because of the lack of appropriate animal models. Modeling ICF syndrome in mice has been discouraging. Complete inactivation of *Dnmt3b* leads to embryonic lethality (8). Although mice carrying ICF-like *Dnmt3b* missense mutations are viable and exhibit some features of ICF syndrome, such as hypomethylation of satellite DNA and cranial facial abnormalities, they do not exhibit antibody deficiency (45). Recent work suggests that Zbtb24 and Hells are involved in class switching recombination (CSR) (46, 47). However, the significance of CSR impairment in immunodeficiency in ICF syndrome remains to be determined. B cells from ICF patients have been shown to be competent for CSR in vitro (20). Many ICF patients show deficiency in IgM, among other Ig isotypes, which cannot be explained by defective CSR. In this study, we show that mice deficient for *Zbtb24* in the hematopoietic lineage (VAV-CKO mice) recapitulate major clinical features of ICF syndrome, including decreased levels of IgM, IgG1 and IgA, thus providing a valuable mouse model of this rare human disease.

Similar to ICF patients, who usually have normal T and B cell counts but lack plasma cells (1,2, 20), VAV-CKO mice exhibit no abnormalities in lymphocyte development but display decreases in plasma cells, suggesting impaired B cell activation and/or terminal differentiation. The immune defects associated with *Zbtb24* deficiency appear to be independent of T cells, as cytokine secretion by helper T cells from VAV-CKO mice is unaltered and, moreover, mice with *Zbtb24* being deleted in T cells (mediated by Lck-Cre) exhibit no obvious immune phenotype. Therefore, our work focused on B cells. We show that MZ B cells from VAV-CKO mice are severely defective in proliferation after in vitro stimulation with the soluble F(ab’)2 fragments of anti-IgM, while their response to LPS stimulation is unaffected. In agreement with the in vitro results, immunization experiments reveal that VAV-CKO mice are hyporesponsive to the TI-2 antigens NP-Ficoll and NP-Dextran but exhibit a normal response to the TI-1 antigen NP-LPS. Antigen-activated MZ B cells rapidly differentiate into plasma cells that secrete antibodies, primarily IgM, that provide innate-like immunity against blood-borne pathogens (48). Our results indicate that activation of *Zbtb24*-deficient MZ B cells by TI-2 antigens is defective, which likely contributes to IgM deficiency in VAV-CKO mice. The decreased levels of other isotypes (e.g. IgG1 and IgA) in these mice could be secondary to IgM deficiency and/or due to CSR impairment (47).

In contrast to diminished TI-2 responses, VAV-CKO mice are hyperresponsive to immunization with the TD antigen NP-KLH. Interestingly, B cell-specific deletion of *Dnmt3a* and *Dnmt3b* in mice results in a similar phenotype: normal B cell development and enhanced TD responses upon immunization (35). Notably, B cells from ICF patients have also been shown to respond more robustly to BCR stimulation than those from healthy donors (20). Taken together, these observations indicate that abnormally enhanced TD responses, mediated mainly by FO B cells, may also contribute to immunodeficiency in ICF syndrome. Our data show that VAV-CKO mice tend to produce low-affinity antibodies in response to immunization, raising the possible involvement of Zbtb24 in antibody affinity maturation.

The divergent responses of VAV-CKO mice to different antigens, i.e. enhanced TD and diminished TI-2 responses, are reminiscent of the phenotype of hCD19TG mice (27). CD19 functions as a thresholding controller of BCR signaling in response to TD and TI antigens, whereby loss of CD19 inhibits TD and TI-I antigen responses in mice while overexpressing hCD19 in mice inhibits TI-II antigen response. We show that, although CD19 expression is not affected by *Zbtb24* deficiency, B cells from VAV-CKO mice, as well as *Zbtb24*-deficeint CH12F3 cells, display substantially elevated CD19 phosphorylation, both before and after stimulation with anti-IgM. Importantly, the immune defects of VAV-CKO mice, including antibody deficiency, can be reverted by disrupting one copy of *Cd19* in these mice. These results demonstrate that Zbtb24 is a negative regulator of CD19 activity and that enhanced CD19 activation is largely responsible for immunodeficiency in VAV-CKO mice. CD19 positively modulates BCR signaling by lowering thresholds for activation and enhancing signaling events (28). Indeed, B cells from ICF patients, relative to those from healthy individuals, are more readily activated in response to BCR stimulation in vitro (20), which may be related to enhanced CD19 activation.

RNA-Seq analysis of naive B cells from VAV-CKO and control mice identified only dozens of differentially expressed genes, including several known Zbtb24 target genes such as *Cdca7, Taf6, Ostc* and *Cdc40* (16, 17), suggesting that Zbtb24 regulates the expression of a limited number of genes. By gain- and loss-of-function experiments using CH12F3 cells, we validated that upregulation of *Il5ra* contributes to enhanced CD19 activation in *Zbtb24*-deficient B cells. Further studies are required to elucidate the mechanisms by which canonical IL-5 signaling and/or non-canonical IL-5Rα functions facilitate CD19 phosphorylation. While the IL-5Rα chain is constitutively expressed on and required for the development and functions of eosinophils and B-1 cells, it is not highly expressed in most conventional B-2 (MZ and FO B) cells (49–53). Indeed, previous work has shown that IL-5Rα, CD19 and other IgM-BCR components (CD79a, CD79b, CD38) are repressed after IgM-BCR stimulation (54). Our results suggest that Zbtb24 is an important factor required for repressing *Il5a* in B-2 cells. Unlike most ZBTB family members, which function as transcriptional repressors (55, 56), Zbtb24 has been associated with gene activation (16, 17, 19). Thus, *Il5a* may not be a direct target of Zbtb24. Given that the Zbtb24-Cdca7 axis regulates the specificity of DNA methylation (19), one possibility is that, in the absence of Zbtb24, *Il5a* is derepressed due to loss of DNA methylation at specific regulatory elements.

In summary, we show that mice deficient for *Zbtb24* in the hematopoietic lineage recapitulate the hypogammaglobulinemia phenotype of ICF patients. Characterization of this animal model reveals that *Zbtb24* deficiency in B-2 cells results in derepression of *Il5a* and elevated IL-5 signaling, leading to enhanced CD19 phosphorylation, which contributes to immunodeficiency, likely by modulating BCR signaling. Therapeutics targeting IL-5, IL-5Rα, CD19, and BCR signaling have been developed for various diseases (57–59). Our findings raise the possibility of using these therapeutics to treat patients with ICF syndrome.

## MATERIALS AND METHODS

### Mice

All animal experiments were performed under protocols approved by the Institutional Animal Care and Use Committee at The University of Texas MD Anderson Cancer Center. The *Zbtb24* conditional allele (*Zbtb24^2lox^*) was generated by gene targeting using homologous recombination. Briefly, mouse embryonic stem cells (mESCs) were transfected with the targeting vector (Figure S1A) via electroporation and selected with Geneticin (G418). Correctly targeted mESC clones (*Zbtb24^3lox/+^*), identified by Southern blotting (Figure S1B), were injected into blastocysts to obtain chimeric mice, which were then crossed with WT mice to obtain F1 mice with germline transmission of the *Zbtb24^3lox^* allele. Germline deletion of the *IRES-βGeo* selection cassette (flanked by *FRT* sites, Figure S1A), resulting in the *Zbtb24^2lox^* allele (Figure S1C), was achieved by crossing *Zbtb24^3lox/+^* mice with FLPeR mice, which constitutively express the FLPe recombinase (60). *Zbtb24^2lox/+^* mice, initially as C57BL/6-129 hybrids, were backcrossed with C57BL/6 mice for six generations. Deletion of the floxed region, resulting in the null allele (*Zbtb24^-^*), in the germline and hematopoietic, T cell, and B cell lineages was achieved by crossing mice carrying the *Zbtb24^2lox^* allele with Zp3-Cre (61), Vav-Cre (31), Lck-Cre (62), and CD19-Cre (36) mice, respectively. All primers used in the study, including those for genotyping and for generating the *Zbtb24* gene targeting vector, are listed in Table S1.

### Immunization

For TD antigen immunization, 8-10-week-old mice were i.p. injected with 100 μg of NP-KLH (Biosearch Technologies) in saline supplemented with 50% (vol/vol) Alum (Thermo Fisher Scientific) and, if applicable, were boosted 60 days later with NP-KLH in saline. Sera were collected from the mice before immunization (day 0) and at day 14, 28 and 56 after the first exposure and, if applicable, at day 14 after the boost. For TI antigen immunization, 8-10-week-old mice were i.p. injected with 100 μg of NP-LPS (Biosearch Technologies), NP-Ficoll or NP-Dextran in saline. Sera were collected from the mice at day 0, 6 and 14. The titers of NP-specific antibody isotypes in sera were measured by ELISA using plates coated with NP(7)- BSA or NP(20)-BSA (Biosearch Technologies) as described previously (63).

### ELISA

Serum concentrations of total IgG1, IgG2a, IgG2b, IgG3, IgM, and IgA were determined by using the Clonotyping System-HRP (Southern Biotech) with the Mouse Ig Isotype Panel (Southern Biotech) as a positive reference standard according to the manufacturer’s instructions. For the determination of NP-specific antibodies, ELISAs were performed as described previously (63).

### Flow cytometry

The following antibodies/reagents were used for staining: B220–APC-H7, CD3-PE, IgM-PE-CY7, IgD-PE, CD43-PE, CD11b-PERCP-CY5.5, CD4-APC-H7, CD8-PE-CY7, CD23-FITC, CD21/35-APC, CD19-FITC, CD19-Alexa-R600, CD62L-FITC, CD44-PE, CD138-PE, IgG1-APC, GL7-FITC, CD95-PE, and propidium iodide (purchased from BD Bioscience). CellTrace Violet Cell Proliferation Kit was purchased from Thermo Fisher Scientific. Single-cell BM or spleen cell suspensions were prepared and stained (63) for analysis and sorting using either a Fortessa flow cytometer (BD Bioscience) or FACSFusion flow cytometer (BD Bioscience). Further analysis was conducted with FlowJo software (v9) for gating and analysis.

### Immunofluorescence

Mice were sacrificed, and their spleens were quickly removed, embedded in Tissue-Tek O.C.T. COMPOUND (ProSciTech), and snap frozen in liquid nitrogen. Cryosections (20 μm) were air-dried for 30 min and then fixed in acetone for 10 min. The sections were air-dried for another 30 min and rehydrated for 10 min in PBS. After blocking with antibody dilution buffer [2% goat serum (Sigma-Aldrich) and 1% BSA in PBS] for 30 min, sections were incubated with B220-, CD4- or PNA-FITC (Sigma-Aldrich) overnight at 4°C. After being washed, sections were permeabilized with 0.1% Triton X-100 (Sigma-Aldrich) in PBS for 5 min. Following the final washing, sections were mounted with Mounting Reagent with DAPI (Life Technologies). Confocal microscopy was performed with Zeiss LSM880.

### RNA-Seq

Splenic MZ and FO B cells were sorted using mouse MZ and FO B Cell Isolation Kit with LS columns (Miltenyi) according to the manufacturer’s instructions. Total RNA of the sorted B cells was isolated with RNeasy Mini Kit (Qiagen). The libraries for RNA-Seq were sequenced using 2×75 bases paired-end protocol on an Illumina HiSeq 3000 instrument. 60-70 million pairs of reads were generated per sample. The raw RNA-Seq readouts were mapped to the mouse assembly reference genome (GRCm38) using TopHat2 and bowtie2 (64, 65), open-source software tools that align RNA-Seq reads to a reference genome. The raw read counts were generated by using htseq-count from HTSeq package (version 0.6.1p1) (66). The overall mapping rates by reads were 92-96%. Differential gene expression analysis was performed with DESeq2 (a R/Bioconductor package) (67), using adjusted pvalue < 0.05 as the significance cutoff. The RNA-Seq data are available on Gene Expression Omnibus (GEO) database with accession number GSE184699.

### RT-qPCR

Total RNA was isolated with TRIzol (Invitrogen) and treated with RNase-free DNase I (Thermo Fisher Scientific). Reverse transcription (RT) was performed using ProtoScript First Strand cDNA Synthesis kit (New England Biolabs) to generate cDNAs. RT-qPCR was performed using iTaq Universal SYBR Green PCR Master Mix (Bio-Rad) on the ABI 7900 Real-Time PCR system (Thermo Fisher Scientific). The primers sequences are listed in Table S1.

### Western blotting

Cell lysates were prepared in lysis buffer (50 mM Tris-HCl, pH 7.5, 150 mM NaCl, 0.2 mM EDTA, and 0.1% Nonidet P-40) supplemented with protease and phosphatase inhibitor cocktail (Thermo Fisher Scientific). A total of 50–100 μg proteins were resolved by SDS-PAGE and transferred to a polyvinylidene difluoride membrane. The following primary antibodies were used: Zbtb24 (MBL, PM085), CD19 (Cell Signaling Technology, 3574), phospho-CD19 (Tyr531) (Cell Signaling Technology, 3571), β-actin (Sigma-Aldrich, A5441), and HA tag (Cell Signaling Technology, 3724). Primary antibodies were detected with an HRP-conjugated goat anti-rabbit or -mouse secondary antibody (Pierce), or HRP-conjugated goat anti-guinea pig (Southern Biotech), and developed with an ECL signal detection kit (GE Healthcare).

### B cell culture, activation and proliferation

The sorted mouse FO and MZ B cells and the CH12F3 cell line were maintained in RPMI 1640 medium supplemented with 10% FBS and 0.05 mM 2-mercaptoethanol. To trace cell proliferation, FO and MZ B cells were labeled with CellTrace Violet (Thermo Fisher Scientific), then treated with either 10ng/ml F(ab’)_2_ Fragment Goat Anti-Mouse IgM (Jackson ImmunoResearch) or Lipopolysaccharide (Sigma-Aldrich) for 72h and analyzed with flow cytometry. For cell activation, naïve B cells or CH12F3 cells were treated with 10ng/ml F(ab’)_2_ Fragment Goat Anti-Mouse IgM or 10ng/ml mIL-5 (R&D Systems) for 5 or 10 min and then collected for Western blotting or RNA extraction.

### Protein overexpression, mRNA KD and CRISPR gene editing in CH12F3 cells

Lentiviral supernatants were produced by co-transfection of the packaging vectors *pMD2.G* (Addgene, #12259) and *psPAX2* (Addgene, #12260) and *pCDH-EF1-FHC* vector (Addgene, #64874) expressing HA-Crisp3 or HA-IL-5Rα, *pLKO.1 puro* vector (Addgene, #8453) expressing control or *Il5ra* shRNAs, or *pL-CRISPR.EFS.GFP* vector (Addgene, #57818) expressing the *Zbtb24* sgRNA and Cas9 into 293T cells and collected 48 h after transfection. For infection, CH12F3 cells (5×10^5^ cells/ml) were cultured in 2 ml of medium and 200 μl of viral supernatants supplemented with polybrene (0.5μg/ml) for 24 h. For protein overexpression and shRNA-mediated KD, the cells were selected with puromycin (1μg/ml) for a week, and viable cells were sorted by FACS to derive KD cell pools. For gene editing with the CRISPR/Cas9 technology, GFP^+^ cells were sorted 24 h after infection (without selection), and single cells were seeded into a 96-well plate to derive pure clones.

### Statistical analysis

Results were analyzed by using Graphpad Prism and were expressed as the means ±SD. The statistical significance was analyzed by Student’s t test.

## ACKNOWLEDGMENTS

We thank Drs. Margarida Santos and Momoko Yoshimoto-Kobayashi for discussions. The work was supported by grants (1R01AI12140301A1 to TC; CA16672 to CCSG Cores at MD Anderson Cancer Center) from National Institutes of Health, Core Facility Support Award (RP170002 to JS) from Cancer Prevention and Research Institute of Texas, and NGS allowances from the Center for Cancer Epigenetics (CCE) at MD Anderson Cancer Center. ZY received a fellowship from the Sam and Freda Davis Fund. TH received a CCE Scholarship.

## REFERENCES

1. M. M. Hagleitner et al., Clinical spectrum of immunodeficiency, centromeric instability and facial dysmorphism (ICF syndrome). J Med Genet 45, 93–99 (2008).

2. C. M. Weemaes et al., Heterogeneous clinical presentation in ICF syndrome: correlation with underlying gene defects. Eur J Hum Genet 21, 1219–1225 (2013).

3. P. Maraschio, O. Zuffardi, T. Dalla Fior, L. Tiepolo, Immunodeficiency, centromeric heterochromatin instability of chromosomes 1,9, and 16, and facial anomalies: the ICF syndrome. J Med Genet 25, 173–180 (1988).

4. M. Jeanpierre et al., An embryonic-like methylation pattern of classical satellite DNA is observed in ICF syndrome. Human molecular genetics 2, 731–735 (1993).

5. C. M. Tuck-Muller et al., DNA hypomethylation and unusual chromosome instability in cell lines from ICF syndrome patients. Cytogenet Cell Genet 89, 121–128 (2000).

6. G. Velasco et al., Comparative methylome analysis of ICF patients identifies heterochromatin loci that require ZBTB24, CDCA7 and HELLS for their methylated state. Hum Mol Genet 27, 2409–2424 (2018).

7. G. L. Xu et al., Chromosome instability and immunodeficiency syndrome caused by mutations in a DNA methyltransferase gene. Nature 402, 187–191 (1999).

8. M. Okano, D. W. Bell, D. A. Haber, E. Li, DNA methyltransferases Dnmt3a and Dnmt3b are essential for de novo methylation and mammalian development. Cell 99, 247–257 (1999).

9. R. S. Hansen et al., The DNMT3B DNA methyltransferase gene is mutated in the ICF immunodeficiency syndrome. Proceedings of the National Academy of Sciences of the United States of America 96, 14412–14417 (1999).

10. J. C. de Greef et al., Mutations in ZBTB24 are associated with immunodeficiency, centromeric instability, and facial anomalies syndrome type 2. Am J Hum Genet 88, 796–804 (2011).

11. E. Chouery et al., A novel deletion in ZBTB24 in a Lebanese family with immunodeficiency, centromeric instability, and facial anomalies syndrome type 2. Clin Genet 82, 489–493 (2012).

12. H. Nitta et al., Three novel ZBTB24 mutations identified in Japanese and Cape Verdean type 2 ICF syndrome patients. J Hum Genet 58, 455–460 (2013).

13. P. E. Thijssen et al., Mutations in CDCA7 and HELLS cause immunodeficiency-centromeric instability-facial anomalies syndrome. Nat Commun 6, 7870 (2015).

14. H. Wu et al., Converging disease genes in ICF syndrome: ZBTB24 controls expression of CDCA7 in mammals. Hum Mol Genet 25, 4041–4051 (2016).

15. C. Jenness et al., HELLS and CDCA7 comprise a bipartite nucleosome remodeling complex defective in ICF syndrome. Proc Natl Acad Sci U S A 115, E876–E885 (2018).

16. J. J. Thompson et al., ZBTB24 is a transcriptional regulator that coordinates with DNMT3B to control DNA methylation. Nucleic Acids Res 46, 10034–10051 (2018).

17. R. Ren et al., Structural basis of specific DNA binding by the transcription factor ZBTB24. Nucleic Acids Res 47, 8388–8398 (2019).

18. M. Unoki, H. Funabiki, G. Velasco, C. Francastel, H. Sasaki, CDCA7 and HELLS mutations undermine nonhomologous end joining in centromeric instability syndrome. J Clin Invest 129, 78–92 (2019).

19. S. Hardikar et al., The ZBTB24-CDCA7 axis regulates HELLS enrichment at centromeric satellite repeats to facilitate DNA methylation. Protein Cell 11, 214–218 (2020).

20. C. E. Blanco-Betancourt et al., Defective B-cell-negative selection and terminal differentiation in the ICF syndrome. Blood 103, 2683–2690 (2004).

21. M. Kraus, M. B. Alimzhanov, N. Rajewsky, K. Rajewsky, Survival of resting mature B lymphocytes depends on BCR signaling via the Igalpha/beta heterodimer. Cell 117, 787–800 (2004).

22. G. D. Victora, M. C. Nussenzweig, Germinal Centers. Annu Rev Immunol 40, 413–442 (2022).

23. E. Meffre, The establishment of early B cell tolerance in humans: lessons from primary immunodeficiency diseases. Ann N Y Acad Sci 1246, 1–10 (2011).

24. P. Engel et al., Abnormal B lymphocyte development, activation, and differentiation in mice that lack or overexpress the CD19 signal transduction molecule. Immunity 3, 39–50 (1995).

25. R. C. Rickert, K. Rajewsky, J. Roes, Impairment of T-cell-dependent B-cell responses and B-1 cell development in CD19-deficient mice. Nature 376, 352–355 (1995).

26. L. J. Zhou et al., Tissue-specific expression of the human CD19 gene in transgenic mice inhibits antigen-independent B-lymphocyte development. Mol Cell Biol 14, 3884–3894 (1994).

27. S. Sato, D. A. Steeber, T. F. Tedder, The CD19 signal transduction molecule is a response regulator of B-lymphocyte differentiation. Proc Natl Acad Sci U S A 92, 11558–11562 (1995).

28. T. F. Tedder, CD19: a promising B cell target for rheumatoid arthritis. Nat Rev Rheumatol 5, 572–577 (2009).

29. J. C. de Greef et al., Mutations in ZBTB24 are associated with immunodeficiency, centromeric instability, and facial anomalies syndrome type 2. Am J Hum Genet 88, 796–804 (2011).

30. M. Cerbone et al., Immunodeficiency, centromeric instability, facial anomalies (ICF) syndrome, due to ZBTB24 mutations, presenting with large cerebral cyst. Am J Med Genet A 158A, 2043–2046 (2012).

31. P. Georgiades et al., VavCre transgenic mice: a tool for mutagenesis in hematopoietic and endothelial lineages. Genesis 34, 251–256 (2002).

32. F. Kiaee et al., Clinical, Immunologic and Molecular Spectrum of Patients with Immunodeficiency, Centromeric Instability, and Facial Anomalies (ICF) Syndrome: A Systematic Review. Endocr Metab Immune Disord Drug Targets 21, 664–672 (2021).

33. D. Sterlin et al., Genetic, Cellular and Clinical Features of ICF Syndrome: a French National Survey. J Clin Immunol 36, 149–159 (2016).

34. C. Kamae et al., Clinical and Immunological Characterization of ICF Syndrome in Japan. J Clin Immunol 38, 927–937 (2018).

35. B. G. Barwick et al., B cell activation and plasma cell differentiation are inhibited by de novo DNA methylation. Nat Commun 9, 1900 (2018).

36. R. C. Rickert, J. Roes, K. Rajewsky, B lymphocyte-specific, Cre-mediated mutagenesis in mice. Nucleic Acids Res 25, 1317–1318 (1997).

37. P. Pfisterer et al., CRISP-3, a protein with homology to plant defense proteins, is expressed in mouse B cells under the control of Oct2. Mol Cell Biol 16, 6160–6168 (1996).

38. S. Sato et al., IL-5 receptor-mediated tyrosine phosphorylation of SH2ISH3-containing proteins and activation of Bruton’s tyrosine and Janus 2 kinases. J Exp Med 180, 2101–2111 (1994).

39. M. Koike et al., Defective IL-5-receptor-mediated signaling in B cells of X-linked immunodeficient mice. Int Immunol 7, 21–30 (1995).

40. K. Horikawa, K. Takatsu, Interleukin-5 regulates genes involved in B-cell terminal maturation. Immunology 118, 497–508 (2006).

41. A. L. Whillock, T. K. Ybarra, G. A. Bishop, TNF receptor-associated factor 3 restrains B-cell receptor signaling in normal and malignant B cells. J Biol Chem 296, 100465 (2021).

42. M. Hulten, Selective Somatic Pairing and Fragility at 1q12 in a Boy with Common Variable Immuno Deficiency. Clinical Genetics 14, 294–294 (1978).

43. L. Tiepolo et al., Concurrent Instability at Specific Sites of Chromosomes 1,9 and 16 Resulting in Multi-Branched Structures. Clinical Genetics 14, 313–314 (1978).

44. L. Tiepolo et al., Multibranched chromosomes 1, 9, and 16 in a patient with combined IgA and IgE deficiency. Hum Genet 51, 127–137 (1979).

45. Y. Ueda et al., Roles for Dnmt3b in mammalian development: a mouse model for the ICF syndrome. Development 133, 1183–1192 (2006).

46. Y. He et al., Lsh/HELLS is required for B lymphocyte development and immunoglobulin class switch recombination. Proc Natl Acad Sci U S A 117, 20100–20108 (2020).

47. A. Helfricht et al., Loss of ZBTB24 impairs nonhomologous end-joining and class-switch recombination in patients with ICF syndrome. J Exp Med 217(2020).

48. A. Cerutti, M. Cols, I. Puga, Marginal zone B cells: virtues of innate-like antibody-producing lymphocytes. Nat Rev Immunol 13, 118–132 (2013).

49. Y. Hitoshi et al., Distribution of IL-5 receptor-positive B cells. Expression of IL-5 receptor on Ly-1(CD5)+ B cells. J Immunol 144, 4218–4225 (1990).

50. M. Kopf et al., IL-5-deficient mice have a developmental defect in CD5+ B-1 cells and lack eosinophilia but have normal antibody and cytotoxic T cell responses. Immunity 4, 15–24 (1996).

51. B. G. Moon, S. Takaki, K. Miyake, K. Takatsu, The role of IL-5 for mature B-1 cells in homeostatic proliferation, cell survival, and Ig production. J Immunol 172, 6020–6029 (2004).

52. K. Takatsu, H. Nakajima, IL-5 and eosinophilia. Curr Opin Immunol 20, 288–294 (2008).

53. T. Yoshida et al., Defective B-1 cell development and impaired immunity against Angiostrongylus cantonensis in IL-5R alpha-deficient mice. Immunity 4, 483–494 (1996).

54. B. Damdinsuren et al., Single round of antigen receptor signaling programs naive B cells to receive T cell help. Immunity 32, 355–366 (2010).

55. S. U. Lee, T. Maeda, POK/ZBTB proteins: an emerging family of proteins that regulate lymphoid development and function. Immunol Rev 247, 107–119 (2012).

56. O. M. Siggs, B. Beutler, The BTB-ZF transcription factors. Cell Cycle 11, 3358–3369 (2012).

57. T. Anagnostou, I. B. Riaz, S. K. Hashmi, M. H. Murad, S. S. Kenderian, Anti-CD19 chimeric antigen receptor T-cell therapy in acute lymphocytic leukaemia: a systematic review and meta-analysis. Lancet Haematol 7, e816–e826 (2020).

58. J. A. Burger, A. Wiestner, Targeting B cell receptor signalling in cancer: preclinical and clinical advances. Nat Rev Cancer 18, 148–167 (2018).

59. S. Principe et al., Treating severe asthma: Targeting the IL-5 pathway. Clin Exp Allergy 51, 992–1005 (2021).

60. F. W. Farley, P. Soriano, L. S. Steffen, S. M. Dymecki, Widespread recombinase expression using FLPeR (flipper) mice. Genesis 28, 106–110 (2000).

61. M. Lewandoski, K. M. Wassarman, G. R. Martin, Zp3-cre, a transgenic mouse line for the activation or inactivation of loxP-flanked target genes specifically in the female germ line. Curr Biol 7, 148–151 (1997).

62. Q. Wang, J. Strong, N. Killeen, Homeostatic competition among T cells revealed by conditional inactivation of the mouse Cd4 gene. J Exp Med 194, 1721–1730 (2001).

63. Z. Ying et al., Histone Arginine Methylation by PRMT7 Controls Germinal Center Formation via Regulating Bcl6 Transcription. J Immunol 195, 1538–1547 (2015).

64. D. Kim et al., TopHat2: accurate alignment of transcriptomes in the presence of insertions, deletions and gene fusions. Genome Biol 14, R36 (2013).

65. B. Langmead, S. L. Salzberg, Fast gapped-read alignment with Bowtie 2. Nat Methods 9, 357–359 (2012).

66. S. Anders, P. T. Pyl, W. Huber, HTSeq--a Python framework to work with high-throughput sequencing data. Bioinformatics 31, 166–169 (2015).

67. M. I. Love, W. Huber, S. Anders, Moderated estimation of fold change and dispersion for RNA-seq data with DESeq2. Genome Biol 15, 550 (2014).

