## Supplemental Figures 1-6 & Table 1 for "Characterization of a mouse model of ICF syndrome reveals enhanced CD19 activation in inducing hypogammaglobulinemia"

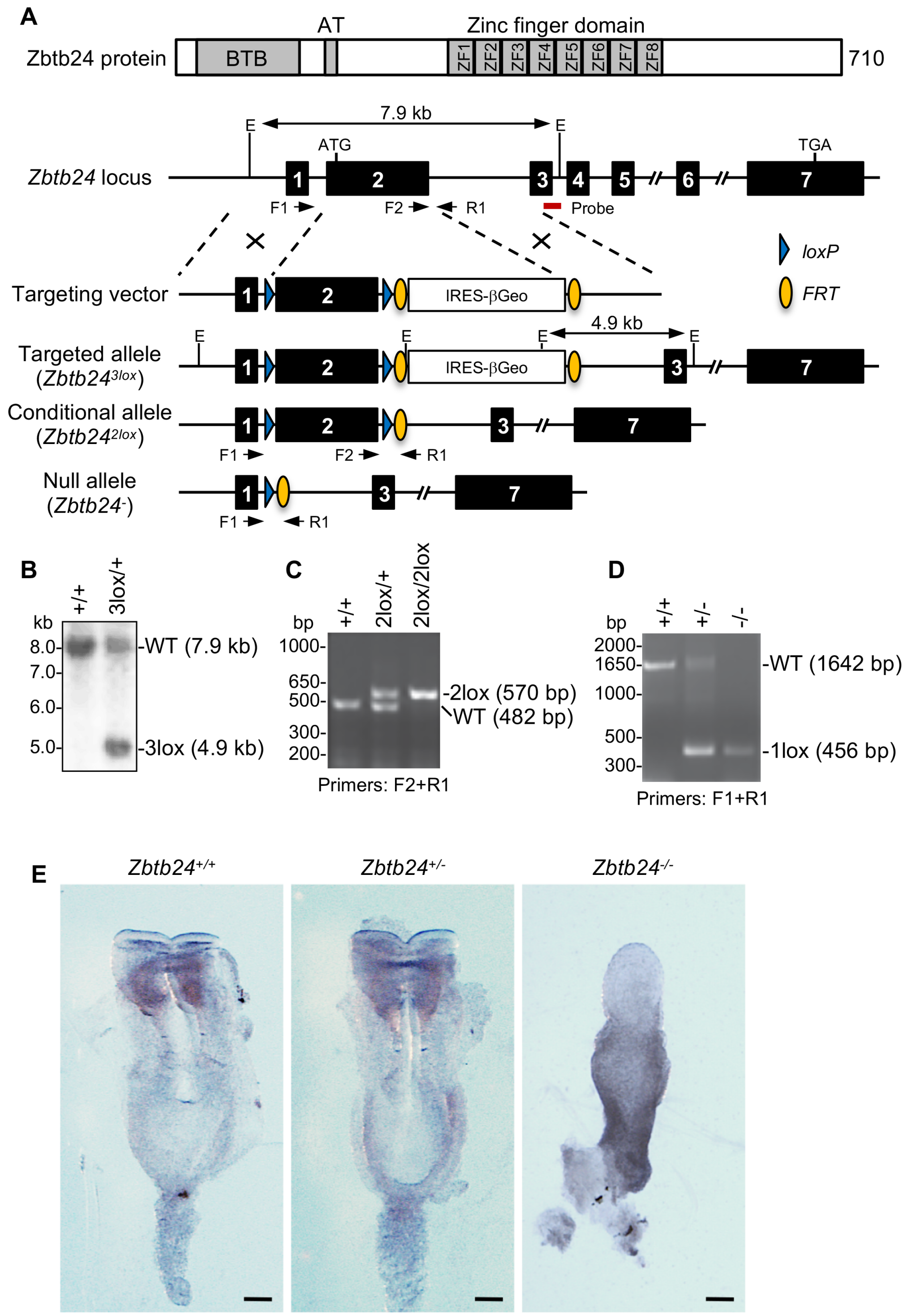


**Figure S1. *Zbtb24* gene targeting strategy and embryonic phenotype of *Zbtb24*-null mice.**
(A) The Zbtb24 protein (including BTB domain, AT-hook, and ZF domain), *Zbtb24* gene locus, conditional KO targeting vector, and various alleles are schematically shown. Exon 2 is flanked by *loxP* sites (blue triangles) in the conditional allele (*Zbtb24*$^{2lox}$) and deleted in the null allele (*Zbtb24*$^{-}$). The initially targeted allele (*Zbtb24*$^{3lox}$) contains an IRES-βGeo selection cassette flanked by FRT sites (yellow ovals). The locations of the PCR primers (F1, F2 and R1) and Southern probe (red bar) used for genotyping, as well as the sizes of diagnostic fragments for various alleles, are indicated. E, EcoRI. (B-D) Representative Southern blot (B) and PCR (C, D) genotyping results. (E) Representative WT (*Zbtb24*$^{+/+}$), heterozygous (*Zbtb24*$^{+/-}$) and homozygous (*Zbtb24*$^{-/-}$) embryos at 8.5 dpc. Scale bars, 100 mm.

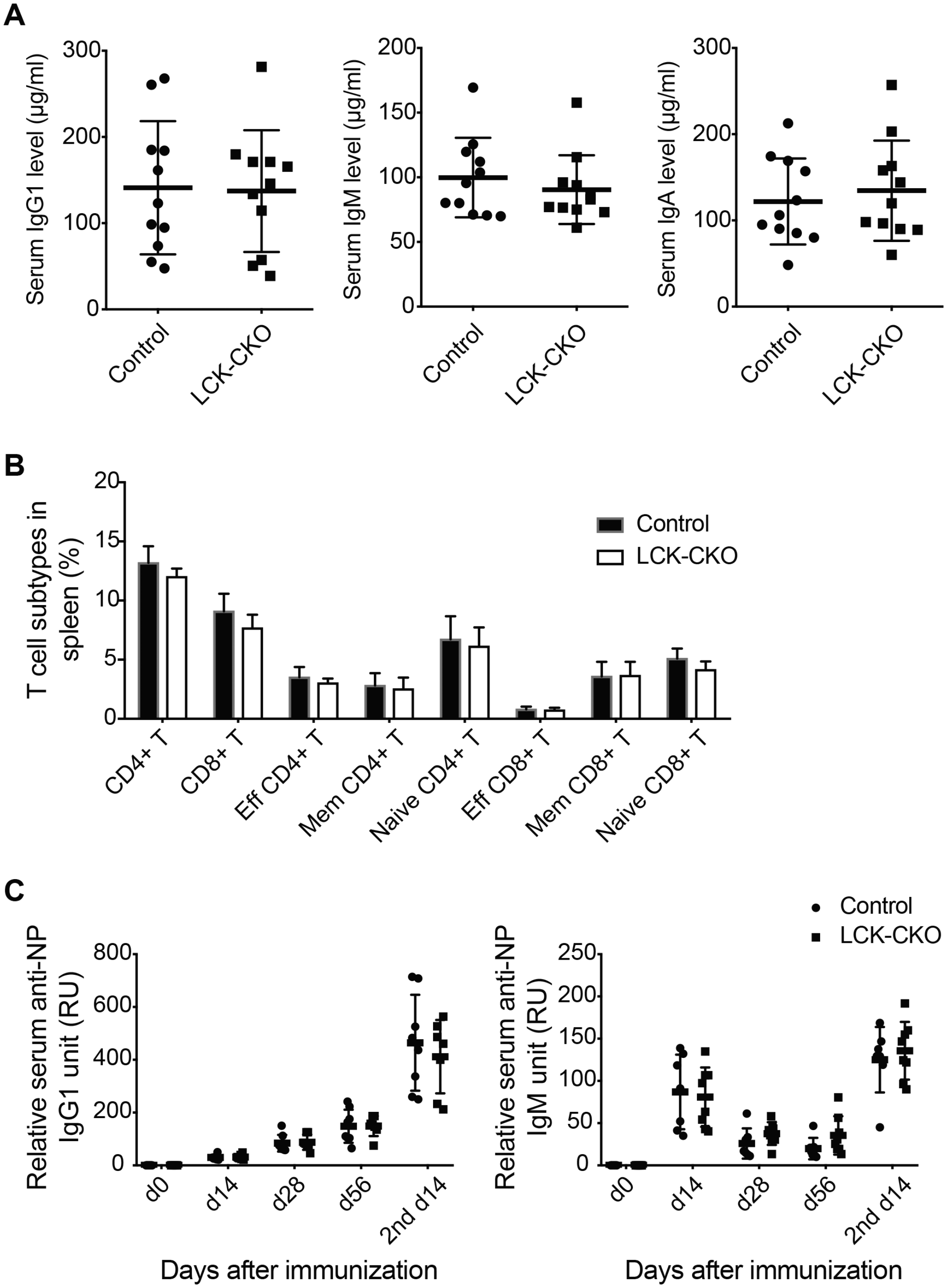


**Figure S2. Lck-Cre-mediated *Zbtb24* ablation in the T cell lineage has no effect on humoral immune response.**

(A) Basal IgG1, IgM and IgA levels in sera from 8-10-week-old Control (*LckCre*:*Zbtb24*$^{2lox/+}$) and LCK-CKO (*LckCre*:*Zbtb24*$^{2lox/2lox}$) mice. (B) Frequencies of T cell subtypes in spleens from Control and LCK-CKO mice. (C) Anti-NP specific IgG1 (left) and IgM (right) absorbance by ELISA in sera from Control and LCK-CKO mice at day 0, day 14, day 56 after first immunization and at day 14 after second immunization with NP-KLH.

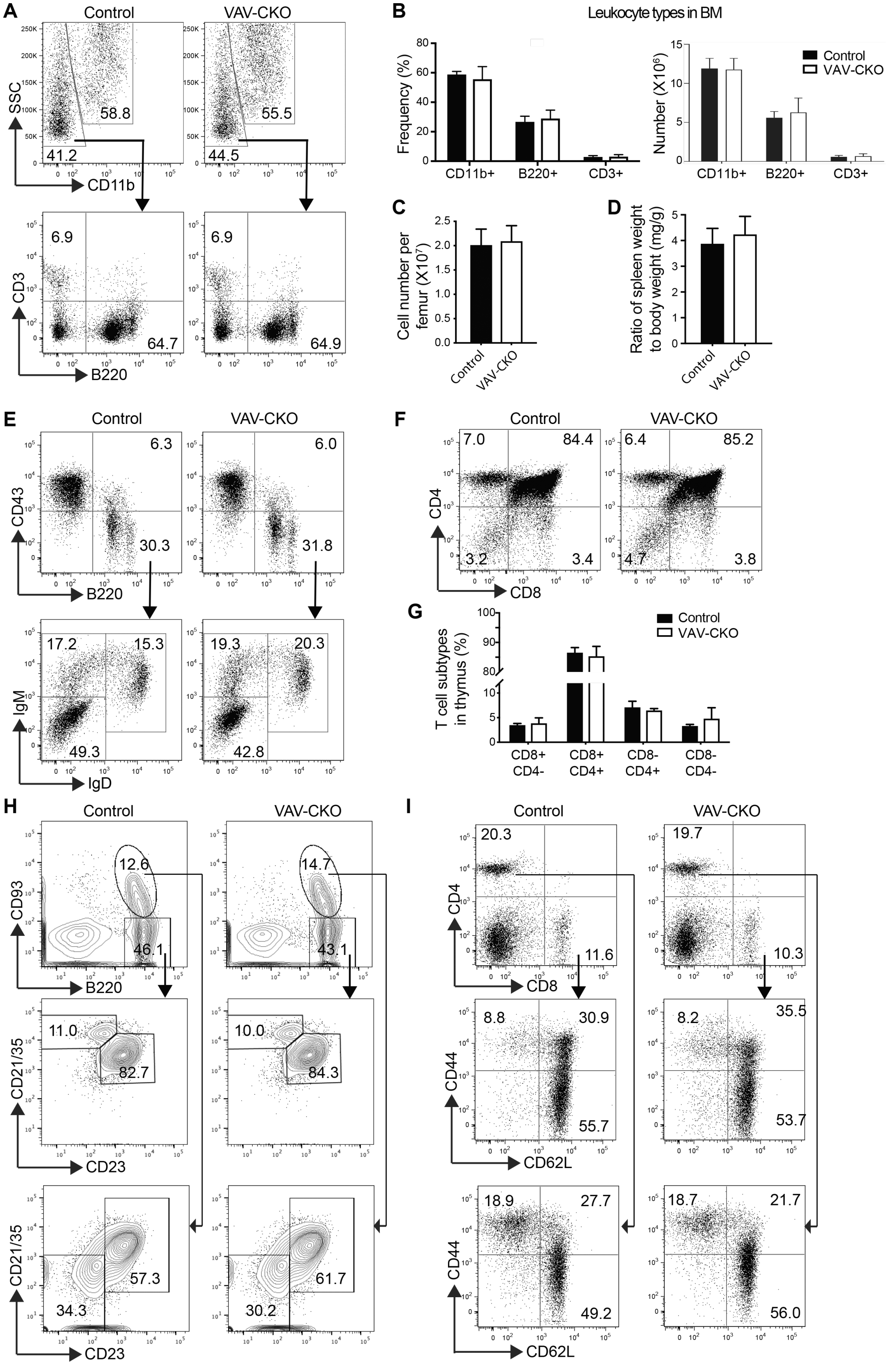
A
Control
VAV-CKO
SSC
CD11b
58.8
41.2
55.5
44.5
CD3
B220
6.9
64.7
6.9
64.9
B
Leukocyte types in BM
Frequency (%)
Number (X10⁶)
CD11b+
B220+
CD3+
Control
VAV-CKO
C
Cell number per femur (X10⁷)
Control
VAV-CKO
D
Ratio of spleen weight to body weight (mg/g)
Control
VAV-CKO
E
Control
VAV-CKO
CD43
B220
6.3
30.3
6.0
31.8
IgM
IgD
17.2
15.3
49.3
19.3
20.3
42.8
F
Control
VAV-CKO
CD4
CD8
7.0
84.4
3.2
3.4
6.4
85.2
4.7
3.8
G
Control
VAV-CKO
T cell subtypes in thymus (%)
CD8+ CD4-
CD8+ CD4+
CD8- CD4+
CD8- CD4-
H
Control
VAV-CKO
CD93
B220
12.6
46.1
14.7
43.1
CD21/35
CD23
11.0
82.7
10.0
84.3
CD21/35
CD23
57.3
34.3
61.7
30.2
I
Control
VAV-CKO
CD4
CD8
20.3
11.6
19.7
10.3
CD44
CD62L
8.8
30.9
55.7
8.2
35.5
53.7
CD44
CD62L
18.9
27.7
49.2
18.7
21.7
56.0

**Figure S3. *Zbtb24* deficiency shows no effect on lymphocyte development.**
(A, B) Flow cytometry analysis of myeloid lineage (CD11b+), T linage (CD3+), and B lineage (B220+) cells in BM from Control and VAV-CKO mice. Shown are representative flow cytometry data (A) and quantification, i.e. frequency (left) and number (right) of the indicated leukocyte cell types (B). (C, D) Total cell numbers per femur (C) and ratio of spleen weight to body weight (mg/g) (D) of Control and VAV-CKO mice. (E) Representative flow cytometry analysis of pro-B (B220+CD43+), pre-B (B220+CD43-IgM-IgD-), immature (B220+CD43-IgM+IgD-) and mature (B220+CD43-IgM+IgD+) B cells in BM from Control and VAV-CKO mice. (F, G) Flow cytometry analysis of T lineage cells in thymus from Control and VAV-CKO mice. Shown are representative flow cytometry data (F) and quantification (G). (H, I) Representative flow cytometry analysis of FO, MZ, T1, and T2/T3 B cells (H) and T cell subtypes (I) in spleens from Control and VAV-CKO mice.

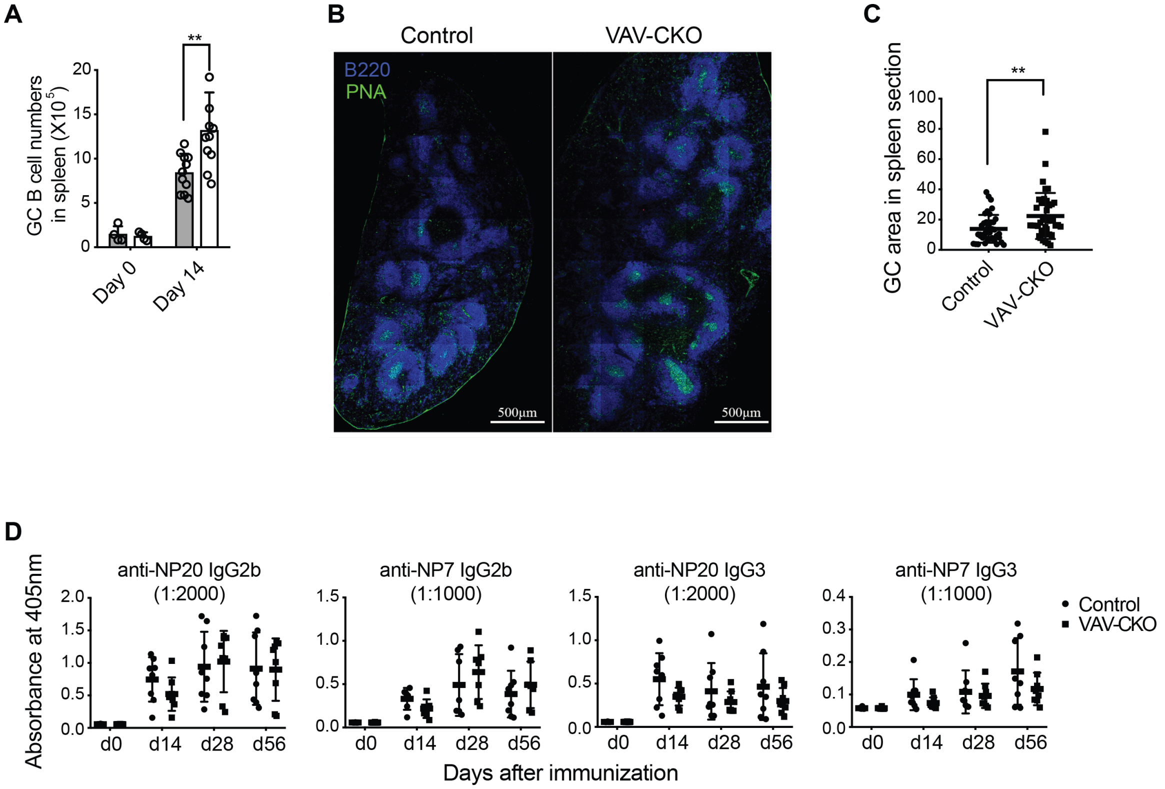


**Figure S4. Immunization with NP-KLH induces enhanced germinal center (GC) B cells in *Zbtb24*-deficient spleen.**

(A) Absolute numbers of GC B cells in spleens from Control and VAV-CKO mice at day 0 and day 14 after immunization with NP-KLH. (B, C) Immunofluorescence analysis of B220 and peanut agglutinin (PNA) in spleen sections of Control and VAV-CKO mice at day 14 after immunization. Shown are representative images (B) and quantification of GC area counts by image J (C). (D) Relative low-affinity (NP20) and high-affinity (NP7) anti-NP specific IgG2b (left two) and IgG3 (right two) antibody absorbance by ELISA in sera from Control and VAV-CKO mice at day 0, day 14, day 28, and day 56 after immunization with NP-KLH.

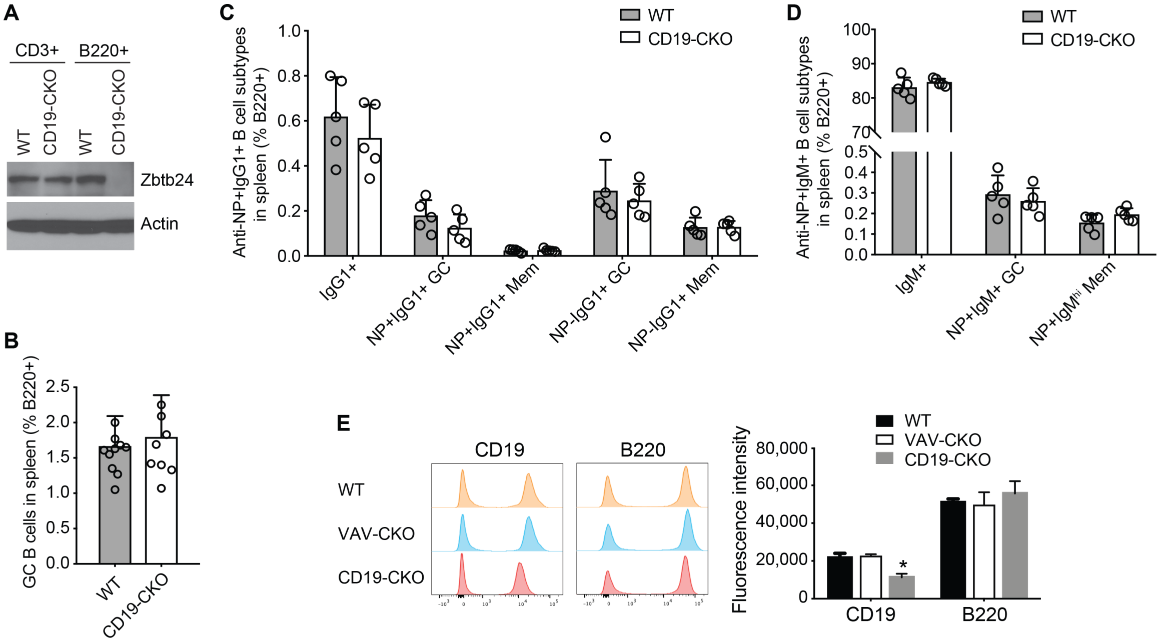


**Figure S5. Characterization of CD19-CKO mice.**
(A) Western blot showing depletion of Zbtb24 in B220+ B cells, but not CD3+ T cells, from CD19-CKO mice. (B) Frequencies of germinal center (GC) B cells at day 14 after immunization with NP-KLH in spleens of WT and CD19-CKO mice. (C, D) Frequencies of anti-NP IgG1+ (C) and IgM+ (D) B cell subtypes in spleens of WT and CD19-CKO mice. Mem, memory. (E) Flow cytometry analysis showing that membrane CD19 is reduced by half on B cells from CD19-CKO mice, as compared with CD19 levels on B cells from WT and VAV-CKO mice. As a control, B220 levels were also analyzed. Shown are representative flow cytometry data (left) and quantification (mean±SD from 3 mice per genotype) (right). Statistical analysis was done using Student's t-test. *$P$<0.05.

**A**

***Zbtb24* Exon 2** — **Targeting sequence** — **PAM**

---GTGGAAGAAAAAAGCGCTCCAGCGGTCCCG**AGG**CCCGGTGTAAAGACTG---

**B**

| Clone No. | Sequence | Indel | Genotype |
| --- | --- | --- | --- |
| 1 | ---GTGGAAGAAAAAAGCGCTCCAGCGGTC-------------TAAAGACTG--- | 13-nt deletion | *Zbtb24*$^{-/-}$ (homozygous) |
| | ---GTGGAAGAAAAAAGCGCTCCAGCGGT**G**CCCGAGGCCCGGTGTAAAGACTG-- | 1-nt insertion | |
| 10 | -AAGATGACC-(78-nt deletion & 29-nt insertion)-TCTTTAAA- | 49-nt deletion | *Zbtb24*$^{-/-}$ (homozygous) |
| | ---GTGGAAGAAAAAAGCGCTCCAGCGGTCC**T**-AGGCCCGGTGTAAAGACTG--- | 1-nt deletion & 1-nt mutation | |
| 11 | ---GTGGAAGAAAAAAGCGCTCCAGCGGTC-CGAGGCCCGGTGTAAAGACTG--- | 1-nt deletion on both alleles | *Zbtb24*$^{-/-}$ (homozygous) |
| | ---GTGGAAGAAAAAAGCGCTCCAGCGGTC-CGAGGCCCGGTGTAAAGACTG--- | | |
| 12 | ---GTGGAAGAAAAAAGCGCTCCAGCG-------GGCCCGGTGTAAAGACTG--- | 7-nt deletion on both alleles | *Zbtb24*$^{-/-}$ (homozygous) |
|  | ---GTGGAAGAAAAAAGCGCTCCAGCG-------GGCCCGGTGTAAAGACTG--- |  |  |
| 15 | ---GTGGAAGAAAAAAGCGCTCCAGCGG**G**CCC-------GGTGTAAAGACTG--- | 1-nt mutation & 7-nt deletion on both alleles | *Zbtb24*$^{-/-}$ (homozygous) |
|  | ---GTGGAAGAAAAAAGCGCTCCAGCGG**G**CCC-------GGTGTAAAGACTG--- |  |  |
| 20 | ---GTGGAAGAAAAAAGC-------------------CCGGTGTAAAGACTG--- | 19-nt deletion on both alleles | *Zbtb24*$^{-/-}$ (homozygous) |
|  | ---GTGGAAGAAAAAAGC-------------------CCGGTGTAAAGACTG--- |  |  |
| 3 | ---GTGGAAGAAAAAAGCGCTCCAGCGGTCCCGAGGCCCGGTGTAAAGACTG--- | WT | *Zbtb24*$^{+/-}$ (heterozygous) |
| | ---GTGGAAGAAAAAAGCGC**C**TCCAGCGGTCCCGAGGCCCGGTGTAAAGACTG-- | 1-nt insertion | |

**Figure S6. Disruption of *Zbtb24* by CRISPR/Cas9 in CH12F3 cells.**
(A) Strategy for generating *Zbtb24*-mutant CH12F3 cell lines with CRISPR/Cas9 technology. Part of exon 2 is shown, with the targeting sequence underlined and the Protospacer Adjacent Motif (PAM) shown in bold. (B) Sequencing results of six *Zbtb24* null clones (1, 10, 11, 12, 15 and 20) and one heterozygous clone (clone 3) are shown.

### Table S1. Primers used in the study

| Name | Sequence (5' to 3') | Application |
|---|---|---|
| shIL5ra-F1 | CCGGCGCCAATAAGTTCATCTCAAACTCGAGTTTGAGATGAACTTATTGGCGTTTTT | shRNA (shIl5ra-1) |
| shIL5ra-R1 | AATTAAAAACGCCAATAAGTTCATCTCAAACTCGAGTTTGAGATGAACTTATTGGCG | shRNA (shIl5ra-1) |
| shIL5ra-F2 | CCGGGCTGACTTACTTAATCACAAACTCGAGTTTGTGATTAAGTAAGTCAGCTTTTT | shRNA (shIl5ra-2) |
| shIL5ra-R2 | AATTAAAAAGCTGACTTACTTAATCACAAACTCGAGTTTGTGATTAAGTAAGTCAGC | shRNA (shIl5ra-2) |
| shIL5ra-F3 | CCGGCCTATTTATGTGGGAAAGGAACTCGAGTTCCTTTCCCACATAAATAGGTTTTT | shRNA (shIl5ra-3) |
| shIL5ra-R3 | AATTAAAAACCTATTTATGTGGGAAAGGAACTCGAGTTCCTTTCCCACATAAATAGG | shRNA (shIl5ra-3) |
| shIL5ra-F4 | CCGGCCAGAAAGACTGAAAGCAAATCTCGAGATTTGCTTTCAGTCTTTCTGGTTTTT | shRNA (shIl5ra-4) |
| shIL5ra-R4 | AATTAAAAACCAGAAAGACTGAAAGCAAATCTCGAGATTTGCTTTCAGTCTTTCTGG | shRNA (shIl5ra-4) |
| Crisp3-qF1 | ATGTGGAGTTGCTGAATGCC | RT-qPCR (Crisp3) |
| Crisp3-qR1 | TGGCCTGCTTGGATAATGGC | RT-qPCR (Crisp3) |
| Crisp3-qF2 | CCATGTGCCAGTTGTCCTGA | RT-qPCR (Crisp3) |
| Crisp3-qR2 | ACATTGGCATGTAGCTAGGCA | RT-qPCR (Crisp3) |
| IL5ra-qF1 | TGCTGGTTTCCCAGGACATTT | RT-qPCR (Il5ra) |
| IL5ra-qR1 | CACGCTTGCTTGAGCCATTA | RT-qPCR (Il5ra) |
| IL5ra-qF2 | TCCAGAATTCGGCcACCATGGTGCCTGTG | RT-qPCR (Il5ra) |
| IL5ra-qR2 | AAATGGATCCAAACGTGGAATTTCCCATGAC | RT-qPCR (Il5ra) |
| Cd19-qF1 | AGTGATTGTCAATGTCTCAGACC | RT-qPCR (Cd19) |
| Cd19-qR1 | CTCCCCACTATCCTCCACGTT | RT-qPCR (Cd19) |
| Zbtb24-sg-F | CACCGAAAGCGCTCCAGCGGTCCCG | CRISPR (Zbtb24) |
| Zbtb24-sg-R | AAACCGGGACCGCTGGAGCGCTTTC | CRISPR (Zbtb24) |
| Vav-Cre-F | GGTGTTGTAGTTGTCCCCACT (in Vav promoter) | Genotyping (Vav-Cre) |
| Vav-Cre-R | CAGGTTTTGGTGCACAGTCA (in iCre) | Genotyping (Vav-Cre) |
| Cd19-Cre-F | AACCAGTCAACACCCTTCC | Genotyping (Cd19-Cre) |
| Cd19-Cre-R | TCAGCTACACCAGAGACGG | Genotyping (Cd19-Cre) |
| Lck-Cre-F | CAGTCAGGAGCTTGAATCCCACGA | Genotyping (Lck-Cre) |
| Lck-Cre-R | TAATCGCCATCTTCCAGCAG | Genotyping (Lck-Cre) |
| Zbtb24-F1 | CGTTCCCATATTTCTCATATATG | Genotyping (Zbtb24) |
| Zbtb24-F2 | GATGGACAGGAGAGCCAGAG | Genotyping (Zbtb24) |
| Zbtb24-R1 | GCAAAGATGAGGAACAGTCATGAG | Genotyping (Zbtb24) |
| C-249 (F) | CTC<u>GCGGCCGC</u>TGCCCTGTGAGTTTTAGTG (5′ arm) | Zbtb24 CKO vector |
| C-253 (R) | CTC<u>ACTAGT</u>ACCCACATAACCTCTTACG (5′ arm) | Zbtb24 CKO vector |
| C-256 (F) | TCGT<u>GTCGAC</u>CTCACAGTAGGTACTATCTC (3′ arm) | Zbtb24 CKO vector |
| C-257 (R) | GAG<u>GGCCGGCC</u>TCTAAAGGTTTCAGAATG (3′ arm) | Zbtb24 CKO vector |
| C-254 (F) | CAT<u>ACCGGT</u>ACTGGAGAGGCATCCAGC (floxed region) | Zbtb24 CKO vector |
| C-255 (R) | GAC<u>CTCGAG</u>ACCTAAACACAACACGAAGG (floxed region) | Zbtb24 CKO vector |

Restriction sites used for cloning are underlined.
